# Neural representations of visual memory in inferotemporal cortex reveal a generalizable framework for translating between spikes and field potentials

**DOI:** 10.64898/2026.01.03.697516

**Authors:** Catrina M. Hacker, Simon Bohn, Brett L. Foster, Nicole C. Rust

## Abstract

Translating neurophysiological findings requires understanding the relationship between common measures of brain activity in animals (spiking activity) and humans (local field potentials, LFP). Prior work suggests alignment between population spiking and high-gamma activity (HGA, 50-150 Hz) of the LFP; however, this is not always observed and is consequently debated. Here, we show that this alignment and prior failures depend on the underlying coding scheme. First, we examined neural representations of variables related to visual memory in macaque inferotemporal cortex. We found that spikes and HGA were strikingly aligned for variables encoded as population response magnitude (novelty, recency, and memorability). However, a variable encoded as a distributed pattern of activity (category) was misaligned. Next, we showed that these insights generalize across many published studies, accounting for prior alignment successes and failures. These results provide a framework for translating between spikes and LFPs, highlighting the scenarios likely to be fruitful for translation.

## Introduction

A core unsolved challenge in translating neurophysiological findings between animal and human studies is understanding how neural representations align across the different ways in which each measures brain activity. In contemporary animal experiments, electrodes are inserted into the brain to record the spiking activity of hundreds of individual neurons [1–3]. In contrast, while invasive clinical measurements in humans can record small groups of single neurons [4], clinical and ethical constraints most often limit these recordings to a more aggregate measure, the local field potential (LFP) [5, 6]. Both measures serve important roles: spiking activity is essential for advancing our understanding of neural coding [7], while field potentials provide insights into human brain function and guide therapies such as brain machine interfaces and neuromodulation [8, 9]. As such, it’s crucial to establish robust translational bridges between spikes and field potentials.

In support of this need, many studies have focused on the extent to which field potentials reflect underlying spiking activity. Historically, the LFP was viewed as reflecting summed synapto-dendritic activity [10– 12], however, several studies have now established that the amplitude of broadband shifts, particularly at high-frequencies, called high-gamma activity (HGA; 50-150Hz), correlates with the activity of single neurons and local neuronal populations in animals and humans (e.g., [13–16]) as well. While this may appear to suggest a distinct biophysical origin of HGA in the LFP, modelling studies have suggested that shifts in HGA may reflect the synaptic integration of local spiking activity [14], reconciling these accounts. This has led human intracranial researchers to widely adopt HGA as a general proxy for local spiking activity to study cognition in the human brain (reviewed by [17–20]). However, the degree of alignment between spikes and HGA continues to be contested. While some studies show alignment between the neural representations measured in simultaneously recorded spikes and HGA [16, 21–25], others show misalignment [22, 24–29], sometimes even for different task variables in the same neural recordings [22, 24, 25], and it’s unclear why. Proposed reasons for misalignment between spikes and HGA include speculation that spikes and LFPs arise from partially distinct phenomena [30], that the relationship between spikes and LFPs varies across brain regions [31], and that the method of LFP spectral analysis influences alignment [15, 32, 33]. Resolving these inconsistencies is important for both research and clinical end-goals when HGA is used as a translational measure of human brain function.

While previous studies have often aimed to bridge this gap by looking at signal correlations between spikes and field potentials alone, we aimed to extend and reconcile these observations by asking what neural representations are common across the two measures. To answer this, we began by comparing how they reflect variables related to visual memory in inferotemporal cortex as rhesus macaque monkeys performed variations of a single-exposure visual memory task. We found that the neural representations for several variables encoded as population response magnitude (novelty, recency, and memorability) were strikingly well-aligned between spikes and HGA. In fact, image novelty was reflected in HGA even more strongly than in spikes, requiring at least 3-fold less data to achieve matched discriminability between novel and repeated images. In comparison, the spiking neural representation of a variable encoded as a pattern of spikes across the population, object category, was poorly captured by HGA. We reconciled these differences by noting that the local neural coding scheme impacts how aggregation across spikes is reflected in the LFP, preserving magnitude but not pattern coding schemes. We then tested the degree to which this difference generalized beyond visual memory to explain alignments versus misalignments in the literature. Across many other brain regions and functional domains we found that variables encoded as magnitude codes were well-aligned between spikes and LFPs while those encoded in the pattern of spiking were not, thus resolving longstanding discrepancies across many studies. Moving forward, this novel, generalizable framework can help identify and predict when electro-physiological findings from animal research will translate to human brain recordings, and in turn, the most promising targets for clinical applications such as brain computer interfaces.

## Results

We began by investigating neural representations of visual memory by comparing spike and LFP data simultaneously recorded from inferotemporal cortex (ITC) of four rhesus macaques (M1-M4) as they engaged in a single-exposure visual recognition memory task [34]. The core task involved viewing a sequence of images and reporting whether each was novel or repeated by making a saccade to one of two choice targets (Figure 1A). Images appeared exactly twice, once as novel (never having been viewed in this or previous experiments) and a second time as repeated. The monkeys initiated each trial by fixating on a center target, after which an image appeared. Fixation had to remain on the image for a minimum amount of time (M1-M2: 500 ms, M3-M4: 400 ms) before two targets appeared and the monkey indicated their choice by making a saccade up or down.

**Figure 1.**
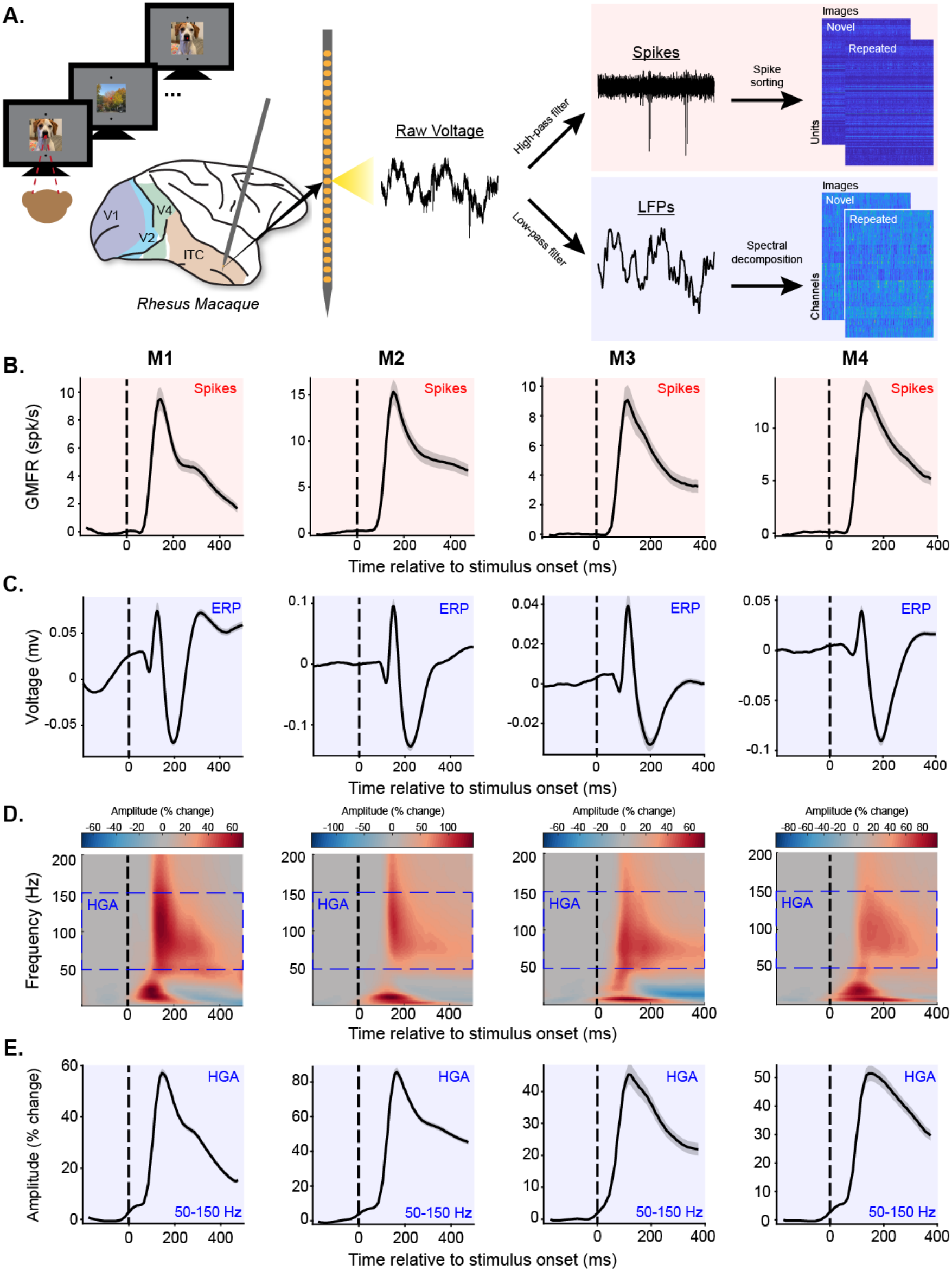
HGA captures population spiking vigor in ITC. Plots highlighted in red show spiking neural activity while plots highlighted in blue show LFP-based neural measures. Each column of B-E contains the data from one monkey (M1-M4) and each row is a different neural measure. **A**. Brain activity was recorded while monkeys performed a single-exposure visual memory task in which they viewed a sequence of images and reported for each whether it was novel or repeated by making a saccade to one of two response targets. While monkeys performed the task, voltage was continuously recorded from inferotemporal cortex of four macaques using a 24-channel probe acutely lowered at the start of each experimental session. Raw voltage was recorded on each channel and either high-pass filtered and manually spike sorted to extract spike times or down-sampled and spectrally decomposed to extract LFP-based measures (see Methods). Data were aligned across sessions to generate pseudopopulations for matched trials with each neural measure (see Methods). **B-E**. Various neural measures as a function of time relative to stimulus onset. Stimulus onset is marked with the dashed vertical black line. Error shadows show bootstrapped 95% confidence intervals across units (spikes) or channels (LFP-derived measures). Several features of the LFP show stimulus-evoked responsivity, but HGA best mimics the spiking grand mean firing rate. **B**. Baseline-subtracted grand mean firing rate as a function of time relative to stimulus onset. Baseline was defined as the mean spiking activity in the window [-200, 0]. **C**. Average voltage fluctuation of the LFP following stimulus onset. **D**. Spectrograms showing percent change in amplitude of each frequency relative to baseline as a function of time relative to stimulus onset. Baseline for each frequency was defined as the mean amplitude in the window [-200, 0]. The blue dashed boxes show the range of frequencies defined as HGA. **E**. Mean HGA as a function of time relative to stimulus onset. In all four monkeys HGA best mimics population spiking vigor (B).

Spike and LFP data were recorded in ITC by acutely lowering a 24-channel U-probe (*Plexon Inc*.) at the beginning of each experimental session. Raw voltage was recorded on each channel and either high-pass filtered and manually spike sorted to extract spike times or downsampled and spectrally decomposed to extract LFP-based measures (Figure 1A, see Methods). Data were aligned across sessions within monkeys to generate pseudopopulations containing simultaneously recorded spike- and LFP-based neural measures for matched trials on their novel and repeated presentations (see Methods). Summaries of statistical tests are reported throughout the text and detailed test statistics can be found in Table S1. The goal of the experiments was to leverage knowledge about the spiking neural representations that support visual memory to systematically compare neural representations measured in spikes and LFPs.

### HGA captures spiking vigor in ITC better than other components of the LFP

Given uncertainty about what conditions facilitate correspondence between spiking activity and the LFP, we began by identifying the features of the LFP that best captured spiking activity. To achieve this, we compared the visually-evoked response (the peri-stimulus time histogram, PSTH) observed in the grand mean firing rate of the spiking population (Figure 1B) with that of different LFP features. First, we examined the average LFP voltage fluctuation in response to image presentation, often called the event-related potential (ERP). The ERP showed consistent responses in all four monkeys (Figure 1C), confirming the presence of visually-evoked signals in the field potential. However, the temporal dynamics of the ERPs did not align well with population spiking (compare Figure 1B and 1C), as quantified by a larger Euclidean distance between the z-scored eventrelated time courses of spiking activity and the ERP (Permutation test (time-shifted surrogates), p>0.05, Table S1). This misalignment was not unexpected given prior work but serves as a valuable reference point for how the ERP, as a neural measure with widespread use in human studies, compares to spiking activity. Given this poor alignment, we sought to identify alternative measures of the LFP that might better capture spiking neural responses.

Following on previous literature suggesting that different frequency bands capture underlying spiking activity to different degrees [15, 16, 22, 25, 27], we next examined how spectral properties of the LFP aligned with underlying spiking activity. Consistent with previous work [16], we found that image presentation was followed, on average, by a characteristic LFP spectral pattern: an early ERP-driven increase in low frequency activity followed (∼80 ms) by an increase in high-frequency activity and decrease in low-frequency activity (Figure 1D). This relationship was observable only after normalizing the amplitude of each frequency with respect to base-line to minimize the effects of the 1/frequency distribution of spectral power. Consequently, all subsequent analyses utilize baseline-normalized spectrograms.

Across canonical frequency bands, previous work has suggested that activity in the high-gamma band (HGA, here defined as 50-150 Hz) aligns more closely with spikes than lower frequency components [15, 16]. Consistent with these observations, we found that increases in HGA induced by image presentation were strong (ranging from a peak of 45.4% to 83.5% increase in amplitude from baseline) and that the temporal dynamics and directionality of HGA tended to align well with underlying spiking activity across all four monkeys, as evidenced by a low Euclidean distance between the z-scored event-related time courses of spiking and HGA (Figure 1E; Permutation test (time-shifted surrogates), p<0.05, Supplemental Table S1). This shared response profile was strongest for HGA compared to other features of the LFP (Supplemental Figure S1) and visually evoked responses were reliably reflected as broadband shifts rather than oscillatory peaks at any frequencies (Supplemental Figure S2). Similarly, given observations that both correlated and asynchronous inputs can be reflected in HGA [11, 35, 36], we ran a series of analyses to examine the relative impacts of the ERP and synchronous spiking activity on HGA (Supplemental Figure S3). While these types of correlated inputs could, in theory, be reflected in HGA, we found that the ERP was reflected in low frequencies but not HGA and that HGA in our data largely reflects asynchronous spiking activity. This is consistent with many previous findings of HGA’s relatively tighter alignment with spiking activity than other frequency bands [14, 20, 37, 38]. Therefore, HGA was our primary focus in the next analyses.

### Novelty signals reflected in spikes and HGA are well-aligned

We consistently observed striking alignment between mean firing rates and mean HGA averaged across all task conditions (Figure 1), so we next sought to investigate how robust this alignment was to task-related modulations. Specifically, we examined how neural representations of several variables relevant to visual memory aligned with HGA. First, we examined the central task manipulation: image novelty. Repeating an image evokes a stimulus-specific reduction in overall population firing rate, a phenomenon known as repetition suppression [34, 39–43]. Repetition suppression is thought to be a neural signal driving memory, as stronger repetition suppression is associated with improved memory performance [34, 44–46]. We observed consistent repetition suppression in population mean spiking activity across all four monkeys (Figure 2A), as evidenced by significantly greater average responses to novel than repeated images in the window from 150 ms following stimulus onset to the end of stimulus presentation (one-tailed Wilcoxon signed rank test, p<0.0001, Supplemental Table S1).

**Figure 2.**
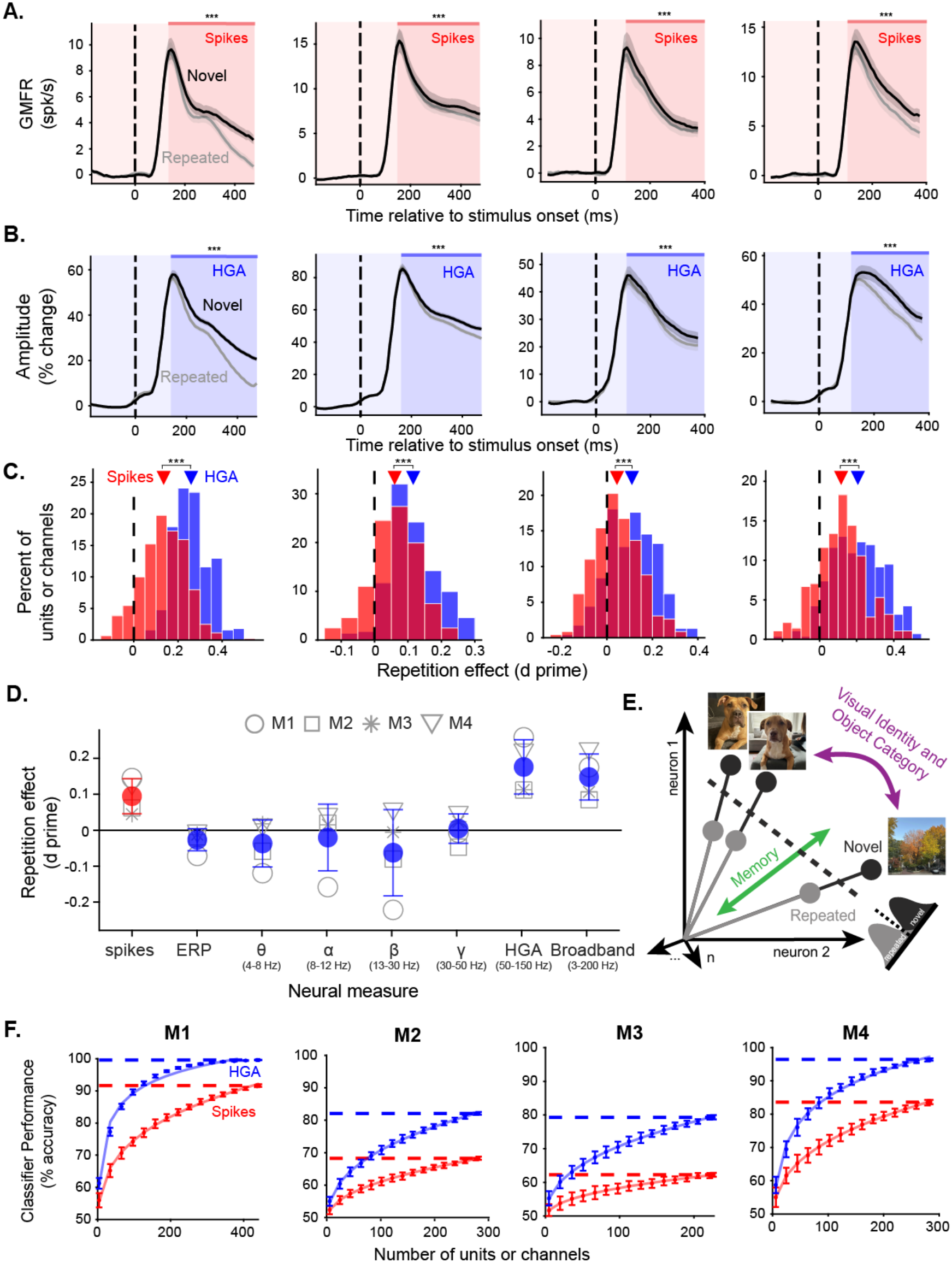
HGA accurately, consistently, and more efficiently captures novelty signals observed in spikes. **A.** Grand mean firing rate across the population as a function of time relative to stimulus onset split between novel (black) and repeated (grey) images for all four monkeys (columns). Shaded area is the window used for significance testing and *** denotes p < 0.0001 for a one-tailed Wilcoxon signed rank test. **B**. Amplitude of HGA (percent change relative to baseline), as a function of time relative to stimulus onset, split between novel (black) and repeated (grey) images for all four monkeys (columns). Shaded area is the window used for significance testing and *** denotes p < 0.0001 for a one-tailed Wilcoxon signed rank test. **C**. Histogram of the repetition effects observed in individual units (red) or channels (blue) measured as a d prime between responses to novel and repeated images for all four monkeys (columns). Positive values correspond to repetition suppression and negative numbers to repetition enhancement. Triangles show the mean for each neural measure. The black dashed line denotes a d prime of 0. *** denotes p < 0.0001 for a two sample t-test comparing the red and blue distributions. **D**. Repetition effect measured as the d prime of responses to novel compared to repeated images for spiking activity (red) and several features of the LFP (blue). D prime was computed separately per unit/channel and then averaged across all units/channels. Open shapes correspond to individual monkeys (circle: M1, Square: M2, asterisk: M3, triangle: M4), filled circles are means across monkeys. Error bars show standard deviation across the four monkeys. **E**. Cartoon depiction of hypothetical ITC responses and the linear classification boundary. Each vector is the population response to one image. Angles between vectors carry information about image identity while the length of the vector denotes the population response magnitude, carrying memory information. The cross-validated linear classifier draws a decision boundary (dashed line) between novel (black) and repeated (grey) images to distinguish them on the basis of their population response magnitude. The cartoon is shown in 2D for simplicity of visualization, but responses are high-dimensional. **F**. Performance of the memory classifier trained on spikes (red) or HGA (blue) as a function of how many units (spikes) or channels (HGA) were included in the analysis shown for all four monkeys. Points are performance and standard deviation across 50 iterations of re-sampling and lines are a power law fit to the data (see Methods).

The repetition suppression signal was also consistently observed in HGA (Figure 2B), as evidenced by greater average responses to novel than repeated images, again considering the window from 150 ms following stimulus onset to the end of stimulus presentation (one-tailed Wilcoxon signed rank test, p<0.0001, Supplemental Table S1). Notably, repetition suppression in HGA captured subtle differences in the spiking activity of individual monkeys, such that the monkey with the greatest repetition suppression in their spikes also showed the greatest repetition suppression in HGA (compare the columns of Figure 2A and Figure 2B), further suggesting that task-related changes in population spiking can be captured by HGA. Repetition suppression in spikes and HGA was also highly consistent, with the majority of individual units and channels showing reduced responses to repeated relative to novel images (Figure 2C). These findings are consistent with prior observations that HGA captures spikes in a small radius around the electrode [47, 48]. Consistent with this interpretation, novelty signals of spikes and HGA on matched channels showed positive but weak correlations (Supplemental Figure S4A).

When comparing across canonical LFP frequency bands, only HGA and broadband activity (3-200 Hz) showed consistent repetition suppression across all four monkeys (Figure 2D, Supplemental Figure S5). Given our decision to normalize our spectrograms relative to baseline, the broadband is dominated by HGA but still includes signals from lower frequencies without repetition suppression, explaining the weaker repetition suppression in broadband activity relative to HGA. The localization of repetition suppression signals to high frequencies was further evident in the full spectrogram of differences between responses to novel and repeated images (Supplemental Figure S4B). Following the stronger novelty signals in HGA, we continued our comparisons between spikes and LFPs by focusing on HGA.

### Novelty signals are stronger in HGA than spikes

While the majority of units and channels showed repetition suppression, individual channels of HGA tended to have stronger repetition suppression than individual units (Figure 2C; two sample t-test, p<0.0001, Supplemental Table S1). To further investigate whether the novelty signals observed in HGA are indeed stronger than those in spikes, we trained a linear classifier to distinguish novel from repeated images and examined how performance scaled with the number of datapoints (spike: units or LFP: channels) included. The cross-validated classifier works by drawing a linear decision boundary to distinguish novel from repeated training images on the basis of their response magnitude (Figure 2E) and classifying an independent test set of images using this decision boundary (see Methods). We interpret classifier performance as a measure of the strength of the novelty signals that can be directly compared between spikes and HGA, as it measures the separability of novel and repeated responses in a cross-validated manner.

The decoding analysis showed novelty signals were consistently stronger across matched numbers of observations in HGA than spikes. For both spikes and HGA classifier performance increased as more units or channels were included in the analysis (Figure 2F). However, performance grew faster for HGA than for spikes, meaning that for matched numbers of observations, responses to novel and repeated images were more separable using HGA than spikes. To capture the rate of growth of classifier performance, we separately fit a power law function for spikes and HGA and compared the estimated amount of data needed to reach 75% classification accuracy (see Methods). Differences ranged from 3.6-to 7.1-fold more data required to reach matched decoding performance with spikes as compared to HGA.

### HGA sensitively captures magnitude modulations related to recency and memorability

The previous analyses focused on novelty signals averaged across all images. However, the single-exposure visual memory task also manipulated two other image-specific variables that impact item-level visual memory: recency and memorability (Figure 3A; described below). Given the presence of strong and consistent novelty signals in HGA that were highly aligned with spikes (Figure 2), we next asked whether the HGA representations were sensitive enough to capture these image-specific neural representations reflected in spikes.

**Figure 3.**
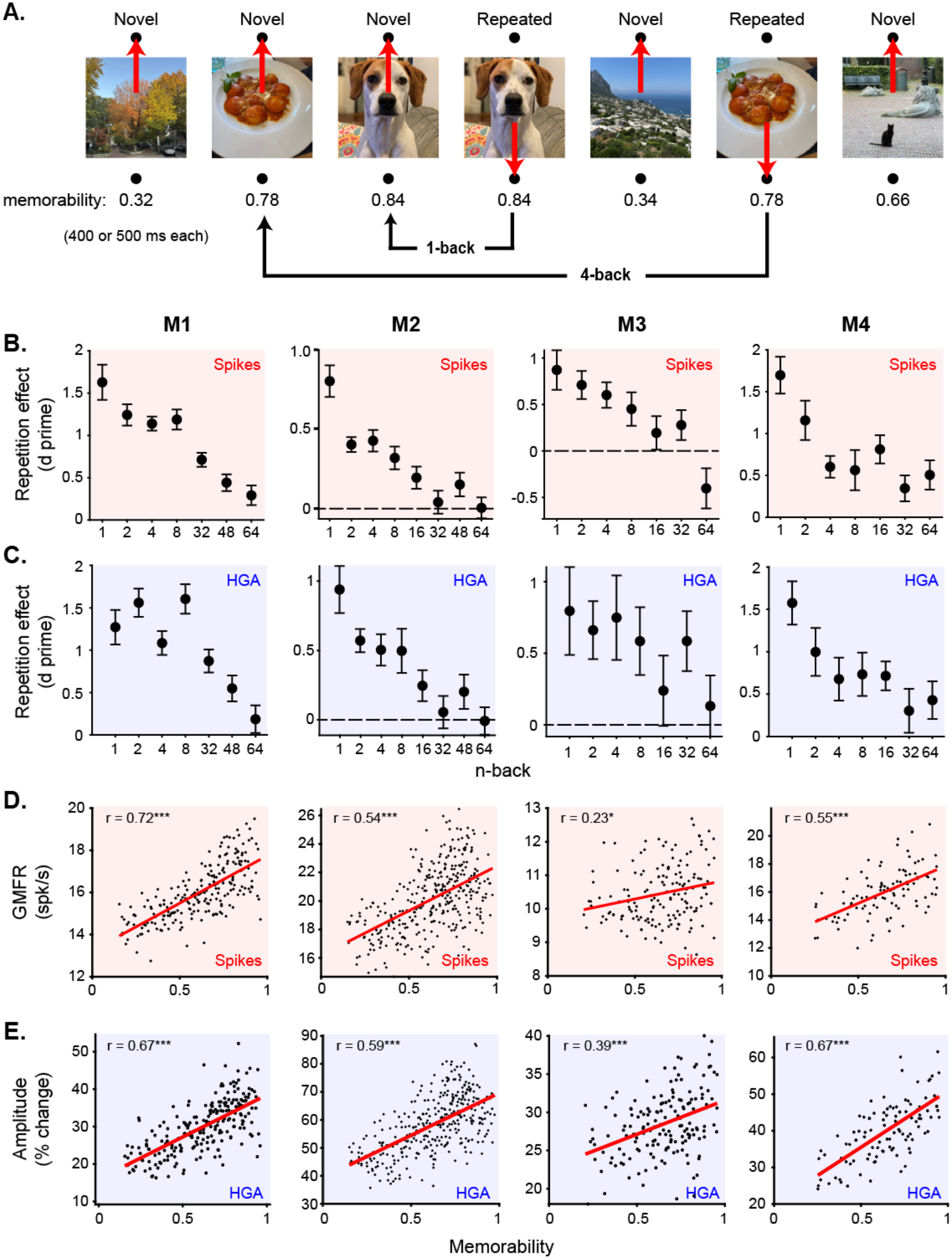
HGA captures image-specific neural representations of recency and memorability. **A.** Illustration of the single-exposure visual memory task. Monkeys viewed a sequence of images and indicated for each whether they thought it was novel or repeated with a saccade up or down (see Methods). Images were presented exactly twice, once as novel and a second time as repeated, with various numbers of intervening images (n-back). Each image was assigned a memorability score using a deep neural network (number in white box; see Methods). **B-C**. Repetition effects (d prime) as a function of n-back for spiking activity (B) and HGA (C) for all four monkeys (columns). Error bars are the standard error of the mean across images. **D-E**. Population grand mean firing rate (D) and HGA (E) as a function of image memorability for all four monkeys (columns). Also shown are a linear regression (red line) and Pearson’s correlation coefficient (r). Statistics for the correlation coefficient are reported as *: p<0.01, **: p<0.001, ***: p<0.0001.

We first considered recency, which was operationalized as the number of intervening trials between the novel and repeated presentation of an image, called n-back (Figure 3A). N-backs ranged from 1 to 64, corresponding to delays of seconds to minutes between image presentations. Consistent with temporal forgetting, memory performance tends to decrease as n-back increases [34] (Supplemental Figure S7). We found that spiking repetition suppression also decreased with n-back (Figure 3B). We thus looked to see whether the same information about recency was aligned between spikes and HGA. We found that HGA repetition suppression was similarly sensitive to recency, systematically decreasing as a function of n-back in all four monkeys (Figure 3C). Importantly, much like in the case of novelty signals (Figure 2A v. B), subtle differences across monkeys in the shape of the relationship between repetition suppression and n-back were consistent across spikes and HGA (compare across columns in Figure 3B and Figure 3C). For example, M2 had the steepest drop-off in repetition suppression as a function of n-back in spikes as well as HGA. This further supports the assertion that HGA sensitively captured underlying spiking activity. We present a comparison of these signals with behavior below, following an assessment of how HGA reflects another task-related variable.

Following our success in capturing recency signals in HGA, which involved averaging responses over several images at matched n-back, we sought to further test the alignment between spikes and HGA by considering representations of image memorability, which required estimating the population response magnitude to individual images in the pseudopopulation. Memorability is an intrinsic image property that captures how consistently memorable an image is across human subjects [49], with values ranging from 0 to 1 denoting least to most memorable images. We found that higher memorability images tended to evoke stronger spiking population response magnitudes in ITC (Figure 3D) (Pearson correlation, p<0.01, Supplemental Table S1). We found that the same was true of HGA, with correlations between image memorability and HGA as strong as with spiking response magnitude (Figure 3E) (Pearson correlation, p<0.0001, Supplemental Table S1). In fact, distributions of estimated response magnitudes of individual images were strikingly similar when computed with spikes and HGA within an individual monkey, as evidenced by strong correlations between population response estimates of spikes and HGA (Pearson correlation, p<0.0001, Supplemental Table S1). Following the same trends as novelty signals (Figure 2D), correlations between response magnitude and memorability were also strongest and better-matched to spikes in HGA than lower-frequency components of the LFP (Supplemental Figure S6).

### HGA predicts behavior as well as spikes using a matched decoding strategy

The correspondence between decreasing repetition suppression (Figure 3B-C) and behavior with increasing n-back suggests a relationship between neural activity and behavior. To more rigorously test this relationship, we asked if a decoder that learns a weighted linear readout to distinguish novel from repeated images on the basis of their response magnitude can predict an individual monkey’s memory performance (see Methods). We trained the same decoder separately on spike or LFP data to test whether the same decoding strategy known to relate spiking activity to behavior [34] can also be applied to LFP data. We focused on recency because, while the mapping between neural representations of memorability and behavior is complex [46], the mapping from recency is straightforward [34].

Cross-validated classifiers were trained to distinguish novel from repeated images and their performance was rescaled to match behavior using methods developed and validated in [34] (see Methods). We found that spikes and HGA were equally good at predicting each monkey’s performance as a function of n-back. Figure 4A shows the behavior and neural predictions when classifiers are trained on spikes or HGA for one example monkey, M4 (other monkeys shown in Supplemental Figure S7). Both spikes and HGA accurately capture the drop off in memory performance with increasing n-back. All other features of the LFP, with the exception of broadband activity (3-200 Hz), gave worse predictions of behavior than HGA (Figure 4B, Supplemental Figure S7) as quantified by comparing a prediction quality (PQ) measure of the match between behavior and decoder predictions (see Methods).

**Figure 4.**
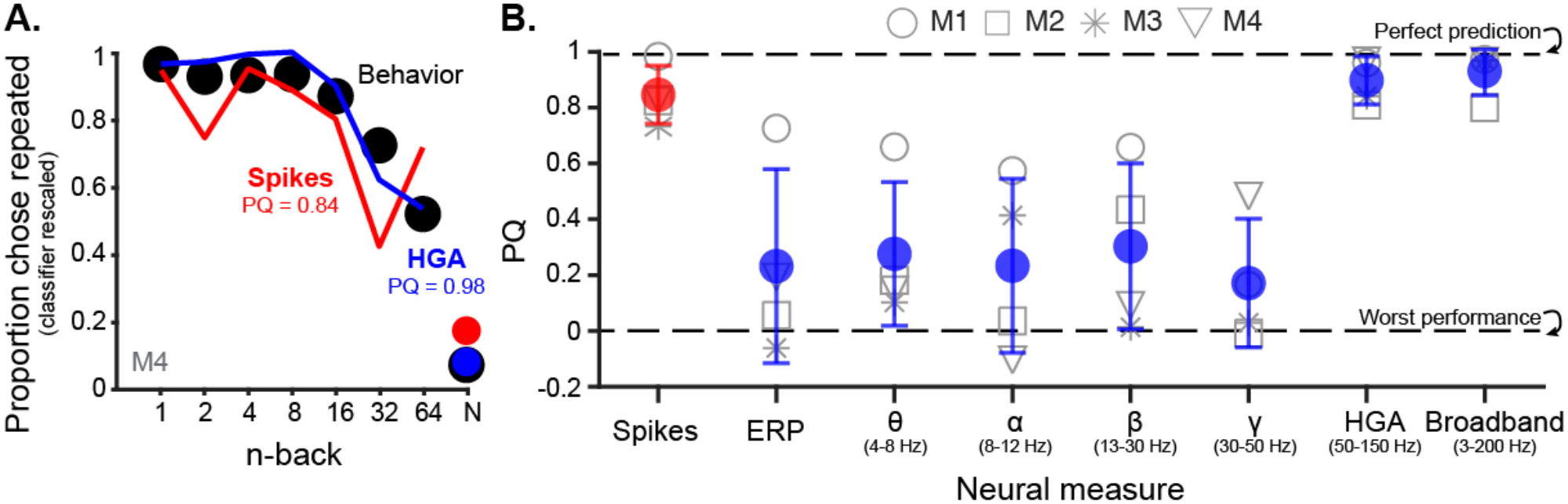
HGA predicts an individual monkey’s behavior as well as spikes using a matched decoding scheme. A. Behavior as a function of n-back (black) and the neural predictions of a classifier trained on spikes (red) or HGA (blue). The x-axis denotes the n-back for repeated images and the average performance for novel images is labeled as “N”. The classifier has been rescaled to match estimated performance for the population size with the lowest MSE (see Methods). **B**. Prediction quality of match between behavior and the classifier predictions trained on spikes (red) and each feature of the LFP (blue) for all four monkeys. The conventions match Figure 2D where each shape is an individual monkey and the colored circles are the means, with error bars showing the standard deviation across monkeys. A PQ of 1 corresponds to perfect prediction, while a PQ of 0 corresponds to chance performance across n-backs (see Methods).

### A pattern-of-spikes code is poorly aligned between spikes and LFPs

Across analyses, we observed strong alignment between neural representations in spikes and HGA, not only to stimulus presentation (Figure 1), but to specific features of visual memory (novelty, recency, and memorability) at the single trial level and consistent with subtle individual differences in spiking responses across monkeys (Figure 2, 3, 4). However, several other studies have reported a much weaker correspondence between spikes and HGA in the encoding of other stimulus variables [22, 24, 26, 27] even within ITC [24, 26, 27]. In considering why our results might be inconsistent with prior literature, we noted that all the variables examined thus far are encoded in spiking activity using the same underlying neural code, which can be distinguished from the population encoding of other stimulus variables. Specifically, we distinguished between two types of neural codes: magnitude codes, which are reflected as correlated, population-wide changes in the overall firing rate of a given brain area and thus change the length of response vectors (Figure 2E, green), and pattern-of-spikes codes, which are encoded as a distributed pattern of spiking activity across a given brain area and thus change the angles of response vectors (Figure 2E, purple). Under the view that HGA reflects a local average of spiking activity, we reasoned that magnitude codes, like novelty, recency, and memorability, might be more robust to spatial averaging and therefore more likely to be reflected in field potentials than pattern-of-spikes codes.

To investigate this hypothesis, we designed a modified version of the single-exposure visual memory task that allowed us to study a pattern-of-spikes code known to be encoded in ITC: object category. The categorical single-exposure visual memory task (Figure 5A) had the same format as the single-exposure visual memory task, but each session contained five blocks of images in which the majority of images (80%) in each block came from a single object category (five categories total per session). While each image was still presented only twice, each category had 64 repeated presentations within a single block (see Methods), providing the power to examine categorical representations. Importantly, we targeted a similar region in ITC and sampled a wide array of object categories so we could investigate a pattern-of-spikes code reflected across a heterogeneous neural population. These populations are distinct from specialized category-selective regions in ITC, such as face patches [50], where increases in local activity across several millimeters of cortex are known to signal specific object categories, which are reliably detected via HGA (e.g. [51]).

**Figure 5.**
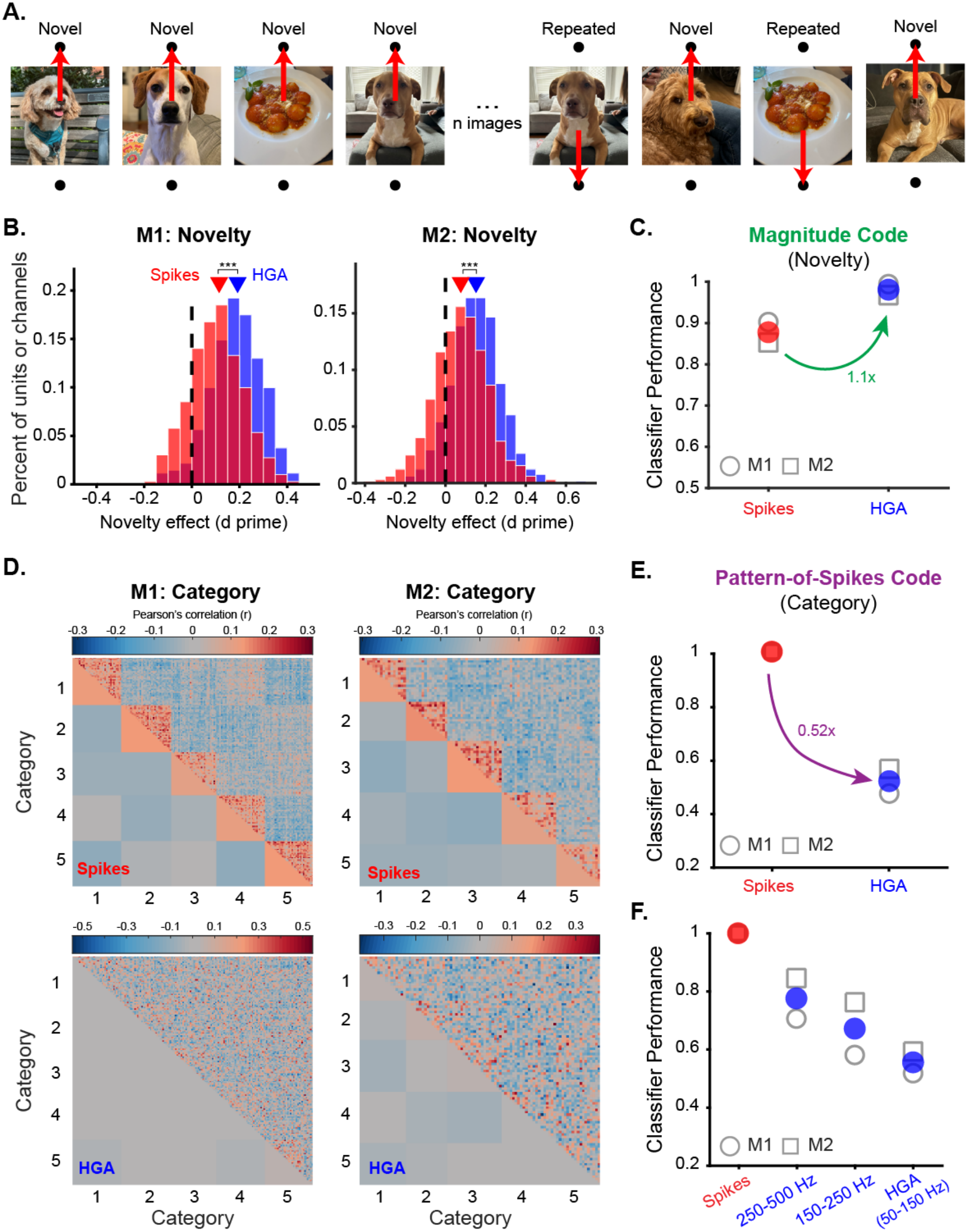
Novelty but not category representations are well-aligned between spikes and HGA. **A.** Illustration of the categorical single-exposure visual memory task. Monkeys viewed a sequence of images and reported whether each was novel or repeated. Trials were organized into blocks where the majority of images (80%) came from the same object category (in this example, dogs). **B**. Histogram of the repetition effects observed in individual units (red) or channels (blue) measured as a d prime between responses to novel and repeated images for both monkeys (columns). Triangles show the mean for each neural measure. The black dashed line denotes a d prime of 0. *** indicates p<0.0001 for a two sample t-test. **C**. Performance of the spike count classifier (see Methods) trained on spikes (red) or HGA (blue) to distinguish novel from repeated images. Open shapes are individual monkeys and colored circles show the mean across the two monkeys. **D**. Representational similarity matrices for neural representations in spikes (top, red box) and HGA (bottom, blue box). Images are grouped into the five categories shown each day. Correlations for individual images are shown above the identity line, and average correlations for each block are shown below the identity line. Color intensity indicates the strength of the correlation. Blocked structure along the identity line illustrates stronger correlations within than across categories, evidence of categorical structure. **E**. Performance of the prototype category decoder (see Methods) trained on spikes (red) or HGA (blue), plotted using the same conventions as C. **F**. Performance of the category classifier for spikes (red) and three frequency bands (blue). Each point is the performance for one monkey and the colored points show the mean across both monkeys. The higher the frequency band the closer neural representations in the LFP match those in spikes, consistent with a decreasing volume of tissue over which neural signals are aggregated.

We first verified that novelty signals in the new dataset were indeed well-aligned between spikes and HGA. We replicated the results in Figure 2C, finding consistent repetition suppression across units (one-sample t-test, p<0.0001, Supplemental Table S1) and channels (one-sample t-test, p<0.0001, Supplemental Table S1) that was stronger in the HGA of individual channels than the spiking activity of individual units (Figure 5B) (two-sample t-test, p<0.0001, Supplemental Table S1). Applying the novelty decoder described in Figure 2F replicated those results, showing strong decoding performance in distinguishing novel from repeated images for both spikes and HGA, but stronger decoding performance with HGA (Figure 5C).

We next assessed categorical representations in spikes and HGA by visualizing the structure of response correlations across all images (representational similarity matrices) produced by each neural measure. Categorical representations should produce blocked structure along the identity line, indicating that the responses to images within the same category are more correlated with each other than they are with responses to images from other categories. The representational similarity matrices produced from population spiking activity did indeed show this blocked structure (Figure 5D, top), indicating strong categorical representations. In contrast, the representational similarity matrices produced from HGA showed a much weaker and nearly absent blocked structure, indicating weak categorical representations (Figure 5D, bottom).

To quantify observed differences in the strength of categorical representations in spikes and HGA in a comparable way to novelty, we trained a simple cross-validated prototype decoder to classify which of the five categories a given image belonged to. On each iteration of cross-validation, one image is selected from each category as the test image and the responses to all remaining images in each category are averaged to produce one “prototype” for each of the five categories. Each test image is classified as belonging to the category whose prototype its response correlates most strongly with (see Methods for details). The classifier results support the observations in Figure 5D, showing ceiling performance when trained with spiking activity and much weaker performance when trained with HGA (Figure 5E). We verified that categorical representations were not decodable in other frequency bands (Supplemental Figure S8A) and that the type of classifier used to decode performance does not qualitatively change the results (Supplemental Figure S8B). We further verified that the anatomically distributed nature of the neural code was what impacted alignment by performing a control analysis in which we demonstrated that HGA and spikes remained poorly aligned even after post-hoc sorting units by their tuning properties (Supplemental Figure S8C).

Altogether, we demonstrate that the representation of a magnitude code (novelty) is much better aligned between spikes and HGA than the representation of a pattern-of-spikes code (category) in the same dataset, suggesting that the nature of the underlying neural code may influence the translation of neural representations between spikes and HGA.

### A generalizable framework accounts for the mapping between spikes and field potentials

Thus far, we’ve shown that the neural representations of several magnitude codes (novelty, recency, and memorability) but not a pattern-of-spikes code (category) are well-aligned between spikes and HGA in ITC. To account for these results, we propose a framework for how neural coding scheme influences neural representations measured in HGA (Figure 6) that builds intuitively on the interpretation that HGA captures a spatial average of local spiking activity. Specifically, when considering HGA, variables that are encoded in population spiking vigor, or correlated changes in population-wide firing rate, are reflected in the length of population response vectors, will average out noise and retain signal, maintaining the relative lengths of response vectors. However, for variables that are encoded as a spatially diffuse pattern of activity across the population, reflected in the angles between response vectors, spatial averaging will average signal away, collapsing population response vectors toward each other (Figure 6, top versus bottom).

**Figure 6.**
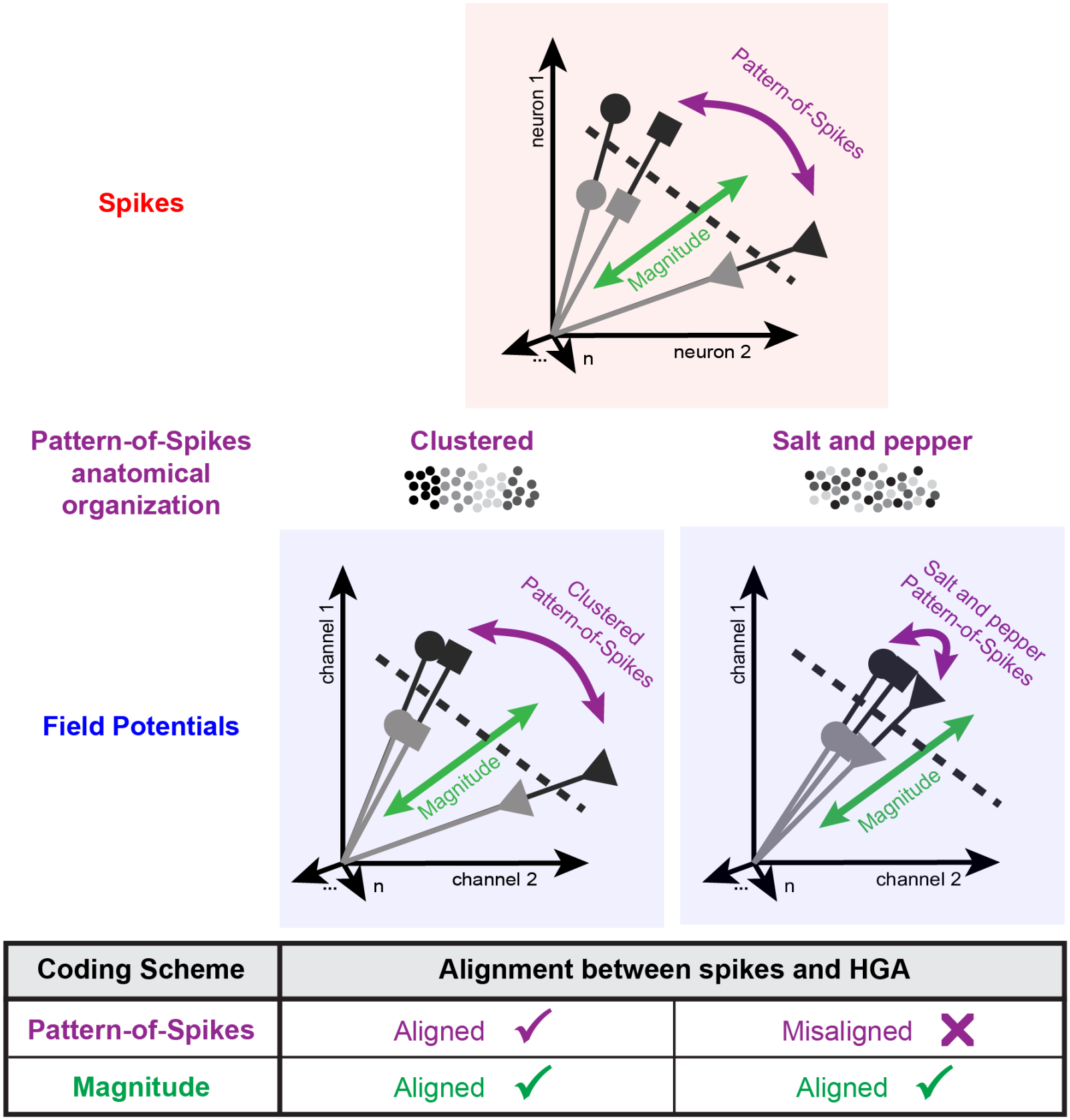
Magnitude and clustered but not salt and pepper pattern-of-spikes codes are aligned between spikes and HGA. Cartoon depiction of neural representations measurable in spikes (top) and field potentials (bottom). Each vector represents a single neural response in high-dimensional space (visualized here in two dimensions) where each dimension is the response of a single neuron (spikes) or channel (field potentials). Dashed lines represent the linear decision boundary distinguishing novel from repeated images. In spiking activity, variables can be encoded as a pattern-of-spikes code (the angles between vectors, purple) or as a magnitude code (the length of the response vectors, green). Compared to representations measurable in spiking activity, the same response vectors measured with field potentials maintain their length, preserving magnitude codes. For pattern-of-spikes codes, however, alignment between spikes and field potentials depends on whether neurons involved in signaling the variable of interest are anatomically clustered or not, with clustered pattern-of-spikes codes maintaining alignment between spikes and field potentials (bottom left), but salt and pepper pattern-of-spikes codes being obscured (bottom right). In the cartoons beneath each label each circle represents a neuron and their color represents tuning for a given variable.

To test our assertion that aggregation of spiking activity in the LFP interacts with coding scheme to impact alignment, we considered the neural representations present in higher frequency bands of the LFP (150-250 Hz, 250-500 Hz, Supplemental Figure S9). Previous work has shown that activity in higher frequency bands tends to converge toward spiking activity[16] and studies that directly estimate the volume of tissue being aggregated in the LFP have found that HGA averages over a larger volume of tissue than higher frequency bands [48, 63]. Therefore, our proposal makes two testable predictions about the neural representations present in higher frequency bands. First, our framework predicts that the neural representations of magnitude codes will remain well-aligned between spikes and higher frequency bands, but that the efficiency of HGA over spikes will decrease as the radius of integration decreases (Figure 2F). The data confirmed this prediction: the neural representations of novelty, recency, and memorability were well aligned between spikes and higher frequency bands (Supplemental Figure S9A-C), and novelty classifier performance was more similar to spikes than to HGA (compare Figure 2F to Supplemental Figure S9E). The second prediction of our framework is that pattern-of-spikes codes will be better aligned between spikes and the LFP at higher frequencies as activity converges toward multi-unit activity. Indeed, we found that category classifier performance increased as frequency band increased, indicating stronger category representations in higher frequency bands (Figure 5F). Together, these observations support our proposal that the averaging of spiking activity over a volume of tissue in the LFP interacts with coding scheme to influence the alignment between spikes and HGA.

Importantly, in our proposed general framework, we distinguish between clustered pattern-of-spikes codes, where neurons signaling the variable of interest are anatomically clustered near each other (Figure 6, bottom left), and salt and pepper pattern-of-spikes codes, in which they are not (Figure 6, bottom right). For example, in ITC the neurons that encode certain categories, like faces, are known to be locally clustered [50], which would constitute a clustered pattern-of-spikes code, whereas the neurons that encode other categories (e.g., refrigerators) are not, and thus constitute a salt and pepper pattern-of-spikes code (explored in Figure 5). We make this distinction because, under the assumption that HGA captures a spatial average of local spiking activity, the tuning of nearby neurons is more likely to be captured by HGA when those neurons have shared tuning profiles than when their tuning is mixed, thus leading to less of a collapse of the population response vector. Together, our proposal suggests that the nature of the underlying neural code in spiking activity can act as a guide, predicting when neural representations will and will not translate between spikes and HGA.

To test the generalizability of this predictive mapping between spikes and HGA, we assessed how well previous studies comparing neural representations in spikes and HGA align with our proposed framework. We collected the results of studies directly comparing neural representations in spikes and field potentials into a table and categorized each variable as being encoded as one of three types of neural population codes: magnitude, clustered pattern-of-spikes, and salt-and-pepper pattern-of-spikes (Figure 6, Table 1). Pattern-of-spikes codes were classified as clustered when neurons involved in signaling the variable of interest were well-established as anatomically clustered near each other.

**Table 1.** Relationship between alignment of neural representations measured with spikes and HGA and the underlying neural code for several stimulus variables. Row colors were arbitrarily assigned to distinguish the three types of neural codes assessed here. Papers were included in the table if they made direct comparisons between neural representations in spikes and field potentials and the reported field potential measures included HGA or broadband activity. Papers that looked for correlations between spikes and HGA with no notion of neural representation were not included in the table. Papers that looked at lower frequency bands, including the gamma band (∼40 Hz), or that did not dissociate between gamma and high-gamma frequencies were not included given previous work suggesting that the gamma band, especially in visual cortex, behaves differently than HGA (e.g., [16, 62]). Variables were classified as aligned if representations in field potentials were captured as well or better than in spikes. Variables were classified as misaligned if the representations measurable in field potentials were weaker than in spikes, including when signals were somewhat aligned, but more weakly than for other stimulus variables (e.g., visual motion direction versus speed in [22]) or the nature of the representations was different even if they could be measured in field potentials (e.g., [28]). The listed studies constitute a diverse sampling of previous studies comparing neural representations in spikes and field potentials for a diverse set of variables and brain regions.

| **Variable** | **Brain Region** | **Aligned?** | **Neural Code** | **Notes** |
|---|---|---|---|---|
| Visual stimulus size[16] | V1 | y | Magnitude[52] | Matched at high frequencies, reversed tuning at low frequencies. |
| Visual stimulus contrast[16, 53] | V1 | y | Magnitude[54] | Matched at high frequencies, reversed tuning at low frequencies. |
| Visual stimulus adaptation[24] | ITC | y | Magnitude[34] |  |
| Somatosensory stimulus intensity[23] | S2 | y | Magnitude[23] |  |
| Motor movement speed[25] | M1 | y | Magnitude[55] |  |
| Visual motion direction[22] | MT | y | Pattern-of-Spikes Cluster[56] |  |
| Saccade direction[21] | LIP | y | Pattern-of-Spikes Cluster[57, 58] |  |
| Visual object identity[27] | ITC | x | Pattern-of-Spikes Salt and Pepper[59] |  |
| Visual object category[26] | ITC | x | Pattern-of-Spikes Salt and Pepper[59] |  |
| Visual motion speed[22] | MT | x | Pattern-of-Spikes Salt and Pepper[60] | Some alignment, but weaker than visual motion direction reported in the same study. |
| Visual shape selectivity[24] | ITC | x | Pattern-of-Spikes Salt and Pepper[59] |  |
| Motor movement direction[25] | M1 | x | Pattern-of-Spikes Salt and Pepper[61] |  |
| Stimulus value[28, 29] | OFC | x | Pattern-of-Spikes Salt and Pepper[28] | Present, but nature is different in LFP |
| Reward value[28] | OFC | x | Pattern-of-Spikes Salt and Pepper[28] | Present, but nature is different in LFP |
| Choice probability[29] | OFC | y/x | Pattern-of-Spikes Salt and Pepper[29] | Single channel better w/HGA, but pooling across channels of HGA is less effective than MUA, consistent with a signal being "washed out" by averaging. |

Consistent with our proposal, across studies we found that the neural representations of variables encoded as magnitude or clustered pattern-of-spikes codes were better-aligned between spikes and HGA than variables encoded as salt and pepper pattern-of-spikes codes (Table 1). This distinction explained differences in alignment across 7 brain areas ranging from visual to prefrontal cortex and 15 task variables ranging from image contrast to reward value across 11 published studies. These findings support the claim that our framework for predicting which neural representations observable in spikes are likely to be captured in HGA may be generalizable across stimulus variables and brain regions.

## Discussion

The ability to invasively record from the human brain has ushered in an exciting new era of neuroscience research [17, 19, 64–69]. However, it has been challenging to connect findings from these human studies with decades of foundational animal research, as they often rely upon different measures of neural activity: spikes and LFPs. Here, we leveraged an understanding of the neural representations of variables related to visual memory in ITC to compare these two measures. Across four monkeys, we found a strong alignment between spikes and HGA for several variables encoded via overall changes in population response magnitude (novelty, recency, and memorability) (Figure 2, 3, 4). In fact, novelty signals were even stronger in HGA than in spikes, requiring at least 3-fold less data to reach matched discriminability of novel from repeated images (Figure 2F). However, representations of a variable encoded as a distributed pattern of spikes (category) were weaker in HGA than in spikes (Figure 5). To explain this difference, we built upon prior proposals that HGA captures an average of nearby spiking activity to propose a novel, generalizable framework in which estimated population vectors collapse toward each other while maintaining their length, obscuring pattern-of-spikes codes and preserving magnitude codes (Figure 6). This framework not only accounted for our results, but a number of long-standing discrepancies across several published studies across a wide diversity of task variables and brain areas (Table 1). Our proposed framework can be used to predict, with sufficient knowledge of the underlying spiking neural representation, when representations are aligned between spikes and field potentials.

In this study, we sought not only to identify correlates of spiking activity in the LFP, but also to clarify what types of functionally significant neural coding schemes may be consistently captured by the LFP. With this goal in mind, we observed consistent alignment between spikes and HGA for visual memory variables (novelty, recency, memorability) encoded as magnitude codes. This strong alignment between spikes and HGA for magnitude codes is broadly consistent with prior studies establishing how the temporal coordination of local spiking activity influences the nature of its aggregation in the LFP and shapes HGA. Event-related activity of local populations (e.g. in response to a stimulus) typically results in temporally coincident but asynchronous firing across cells, which is thought to result in the more broadband increase in spectral power reflected by HGA [14, 20], and accounts for the tight temporal similarity in spike and HGA response profiles. This is distinct from states of highly synchronous and rhythmic firing, which can also be reflected in HGA and result in oscillatory peaks (i.e. gamma oscillations) and spike-field coherence in the LFP spectrum [36]. Consistent with these findings, we found that the broadband shifts in HGA studied here largely reflect asynchronous co-firing of local populations as opposed to correlated changes in electrical activity such as in the ERP or synchronous spiking activity (Supplemental Figure S3). This aligns with prior observations in humans noting that spike-HGA coupling is strongest for neurons that co-fire [13] and a recent study examining the relationship between spikes and HGA (defined as 200-300 Hz) by teaching monkeys to dissociate spiking from HGA recorded on the same shank of a Utah array during a cursor BCI task [38]. In that study, local spiking could be dissociated from HGA on the same shank, while more diffuse neural co-firing was a better predictor of HGA. While we were not testing the biophysical origins of HGA directly, our results are compatible with accounts that HGA captures spiking activity via synaptic integration. Increases in HGA due to asynchronous spiking were captured by filtering spiking activity through a synaptic kernel [14] (Supplemental Figure S3C), consistent with [38]’s finding that HGA correlates best with population cofiring. Further characterizing how HGA may arise from synapto-dendritic currents will be an important area for future research. Where prior studies establish the firing conditions that favor the aggregation of spiking activity into HGA, we build upon them to note how the neural coding scheme interacts with these conditions. Specifically, we find that magnitude coding schemes produce the kind of temporally coincident asynchronous firing of local populations that aligns spiking activity and HGA. When a sufficient number of local neurons fire similarly in response to a common event, the temporally coincident increase in spiking activity results in an increase in HGA amplitude.

We observed that HGA is not only well-aligned with spikes when considering several magnitude codes but even captures novelty signals more strongly than spikes (Figure 2F). This is likely due to how a host of local bio-physical events contribute to the aggregate LFP. While spike sorted data reflects only a subset of local neurons, HGA reflects a more aggregate signal shaped by the contributions from many more local neurons, including influences from non-spiking components of the extracellular field that have also been shown to contribute to the LFP [10–12, 30]. Therefore, while spikes and HGA look highly aligned temporally, they can differ in their information content. The relatively higher strength of novelty signals in HGA than in spikes likely reflects the benefits of the larger “radius of integration” when aggregating responses under conditions where diffuse, large magnitude co-firing occurs, such as repetition suppression in ITC.

Regardless of the specifics of the biophysical origins of HGA, that HGA captures some aspects of spiking activity makes it a practically useful correlate for engineering brain computer interfaces and neuromodulatory devices. One consideration is that the higher information content we observed in HGA is sensitive to electrode size and spacing. The radius of integration is proportional to the size of the recording electrode [48], with larger electrodes aggregating signals over a larger volume of tissue expected to show an even greater advantage to aggregating signal. However, when the radius of integration across recording channels overlaps or the contribution of each spike can be weighted by its informativeness the information content on each electrode becomes correlated and the relative advantage of adding channels will decrease (e.g., as discussed in [70]). Another consideration when designing brain machine interfaces and neuromodulatory devices is that stability and data efficiency are essential (e.g., reviewed in [71]). Our demonstration of the efficiency and specificity of the alignment of HGA to spiking activity, coupled with the ease of calculation, has important implications for the design of these devices. Specifically, we show that when considering variables encoded as magnitude codes it may be better to utilize recording devices with larger contacts specialized to capture aggregate signals in HGA than traditional penetrating microelectrodes, which would be preferable when spiking activity is necessary, such as when considering variables encoded as a pattern-of-spikes code.

While attributes of magnitude codes favor their aggregation and detection in the LFP, pattern-of-spikes coding schemes will hinder aggregation. For example, we observed that categorical representations in spikes were only weakly reflected in HGA. Importantly, our claim is not that LFPs can’t decode category, but that they will fail to do so in areas where category information is represented in a spatially diffuse pattern of spikes, rather than as a local change in magnitude (e.g., clustered versus salt and pepper organization, Figure 6, Table 1). For example, our recordings were not done in canonical category selective regions of ITC (e.g., face and body areas [50, 72]), so while category information could be decoded, it was through a salt and pepper pattern of spiking scheme, which limited alignment with HGA. If these recordings were performed in patches of visual selectivity, such as face patches, local spiking activity would show clear category-specific magnitude changes that are known to be detectable with HGA (e.g., [51]). Indeed, when considering previous publications, we found that pattern-of-spikes codes in which neurons with similar functional tuning are spatially clustered near each other are better aligned between spikes and HGA than those with a salt-and-pepper organization (Figure 6, Table 1). Therefore, our claim is not about which specific representations HGA can detect, but that detection is sensitive to what kind of spiking patterns can be tracked.

Our framework can act as a guide for future researchers interested in translating insights in both directions between spike and field potential data. In moving from spike to field potential data, it can help identify promising targets for translation into clinical applications like brain computer interfaces. In the other direction, it can help interpret insights gleaned from HGA in the absence of spiking activity. HGA has been widely adopted as a proxy for spiking activity in human intracranial work [17–20]. Our results support the use of HGA as a proxy for spiking activity but clarify under what conditions this relationship does and does not hold. Further, our results suggest that observing HGA related to a given variable is predictive of neural coding scheme, in that not all neural coding schemes will produce measurable signals in HGA (Figure 6). Thus, the ability to measure functionally-relevant HGA signals provides hints about spiking neural representations, even when spiking activity cannot be recorded.

This work aims to inform the interpretation of human intracranial recordings and the translation of insights between animal and human recordings. In that light, one limitation of the current work is that the electrodes typically used to record from human patients are much larger than those in the present study [73] and average over a larger area of tissue [48]. A potential concern might be that our findings reflect a more precise LFP recording available with smaller contacts and will not generalize to iEEG recordings which have a larger pooling factor. However, previous work comparing LFPs and ECoG suggests that the nature of the neural signal captured by each method is comparable and that differences between the two recordings are only a matter of scale [48]. Our results suggest that averaging over larger areas of tissue should maintain or increase the signal of magnitude codes like novelty, so long as neural responses remain homogeneous at the increased spatial scale of the larger contact. Similarly, what constitutes a clustered versus salt-and-pepper pattern-of-spikes code (Figure 6, Table 1) must consider a larger area of tissue when considering human iEEG recordings than the LFPs studied here (e.g., the size of motion direction tuning columns that facilitates alignment in [22] may not facilitate alignment with the larger pooling area of larger electrode contacts). Another challenge in comparing our results to human recordings is that there are often many fewer contacts in a given brain region in an individual patient than studied here. Our results suggest that a single observation of HGA can be stronger than for a single neuron when considering a magnitude code (Figure 2F), but a limited number of observations remains an additional challenge of analyzing human compared to animal recordings.

Our study establishes that HGA sensitively and efficiently captures visual memory representations in ITC and variables reflected as magnitude codes. Magnitude codes are abundant across the brain [74] and used to reflect variables including evidence accumulation [75], attention [76] and mood [77]. That magnitude codes can be efficiently captured in HGA suggests they are a productive touchpoint for translation between animal and human neuroscience. Efforts to identify general principles that can facilitate translation between animal and human neuroscience like those reported here will play an important role moving forward in the effort to bridge basic animal research with human clinical applications.

## Acknowledgements

This work was supported by the National Eye Institute of the NIH (award R01EY020851 to N.C.R., award R01MH139654 to B.L.F., and T32EY007035 to support C.M.H.), the National Science Foundation (award 2043255 to N.C.R) and the Simons Foundation (Simons Collaboration on the Global Brain award 543033 to N.C.R.).

## Methods

Experiments were performed on four adult rhesus macaque monkeys (*Macaca mulatta*) with implanted head posts and recording chambers. M1, M3, and M4 were male and M2 was female. All procedures were performed in accordance with the guidelines of the University of Pennsylvania Institutional Animal Care and Use Committee.

The goal of this paper was to leverage previously reported insights about the relationship between spiking activity and neural representations of visual memory to study the relationship between neural representations in spikes and field potentials. As such, data from three sources are included in this paper. The first includes behavioral and neural data from M3 and M4 performing the single-exposure visual memory task. Experimental procedures and spiking data from this source have previously been reported in [34] and [78]. Those reports focused on spiking neural representations of novelty, recency, and memorability (replicated here in Figure 2A, Figure 3B, Figure 3D, and Figure 4). This report meaningfully extends on these findings to determine how those representations align with unpublished field potential data, which cannot be inferred from previous reports. The second source includes behavioral and neural data from M1 and M2 performing the single-exposure visual memory task. Experimental procedures and spiking data from this source have been reported in [46]. That report focused on how spiking neural representations of memorability in ITC can be decoded, and included the results reported in Figure 3D. This report utilizes that dataset to replicate the findings previously reported in [34] for M3 and M4 and to determine how they align with unpublished field potential data, which is not addressed in and cannot be inferred from the previous reports. Finally, the third source is from M1 and M2 performing the categorical single-exposure visual memory task whose spike and field potential data are reported here for the first time.

### Single-exposure visual memory task

Monkeys were trained using standard operant conditioning (juice reward) with head stabilization. Custom software (*The MWorks Project*) was used to display images on an LCD monitor and to collect behavioral responses. Eye movements were tracked using high-accuracy, infrared video eye tracking (Eyelink 1000, *SR Research*).

The core task was to view a sequence of images, one at a time, and to identify for each whether it was novel or repeated. Images were presented exactly twice: once as novel (never having been seen on this or prior days) and a second time as repeated (having been seen exactly once prior). Each trial was self-initiated by fixating on a red square (0.25°) on the center of a gray screen. After 200 (M3 and M4) or 500 (M1 and M2) ms a 4° stimulus appeared. The monkeys maintained fixation on the image for 400 (M3 and M4) or 500 (M1 and M2) ms, at which time the red square turned green (go cue) and the monkey made a saccade to a target 8°above or below the image to indicate whether the image was novel or repeated. The mapping between up and down and novel and repeated was balanced across monkeys so that two made a saccade up for novel and the other two for repeated. For M3 response targets appeared with stimulus onset and for M1, M2, and M4 response targets appeared with the go cue. For M3 and M4 the image stayed on the screen until a fixation break was detected and for M1 and M2 the image disappeared from the screen when the response targets appeared. Images were drawn from the image set whose acquisition was previously reported in [34]. Images used during experimental sessions were not used for training in the same monkey.

The sequence of images presented each day was randomly generated using a custom Matlab script so that each image appeared exactly twice. The script filled set numbers of trials at specified n-backs and then filled a small number of trials at off-target n-backs to complete the sequence. Due to differences in overall behavioral performance, the distributions of n-backs were slightly different for all four monkeys, but all captured a range of n-backs spanning from seconds to minutes. M1 was presented with n-back trials of 1, 2, 4, 8, 32, 48, 64, and 192. The distribution was uniform for n-backs between 2 and 64, while 1- and 192-back trials were half as frequent. 192-back trials were included as catch trials and were not included in these analyses. M2 was presented with n-back trials of 1, 2, 4, 8, 16, 32, 48, and 64, with a uniform distribution between 2 and 48 and half-frequencies for 1 and 64. M3 and M4 were presented with n-back trials of 1, 2, 4, 8, 16, 32, and 64 with a uniform distribution.

Behavior as a function of n-back (Figure 4A, Supplemental Figure S7) was computed as the mean proportion of trials each monkey selected the “repeated” target across all trials and all sessions included in the neural analysis (see Neural Recording).

### Categorical single-exposure visual memory task

The categorical single-exposure visual memory task was recorded using the same experimental set up, trial timing, and image display methods as the single-exposure visual memory task for M1 and M2. Rather than presenting images from random categories throughout the day, the categorical version of the task consisted of seven blocks of 160 trials. Five were categorical blocks wherein 80% of the images were sampled from the same image category (Figure 5A) and 20% were from random image categories. The other two blocks were random blocks where all images came from random categories. Only the responses to categorical images are analyzed here. Images were shown exactly twice and always in the same block. Images were sampled from Ecoset [79], cropped to a square, and resized to 256*256 pixels.

Unlike the single-exposure visual memory task, the image sequences for the categorical task were enriched for a small range of target n-backs (M1: 33-36, M2: 7-10). Target n-backs were chosen for each monkey to keep behavioral performance approximately matched. In addition to the enriched n-backs, several 1-, 2-, and 4-back trials were shown throughout the task as well as a small number of trials at other n-backs used to fill the sequence. Only the trials shown at the target n-backs are included in the analysis.

### Neural recording

Data were recorded in ITC of all four monkeys via a single recording chamber (*Crist Instruments*) whose placement was guided by anatomical magnetic resonance imaging. The region of ITC recorded was located on the ventral surface of the brain, over an area that centered 17mm lateral to the sagittal plane defined by the midline and 13-17 mm anterior to the ear canals, targeting anterior ITC. Neural signals were recorded with a linear 24-channel U-probe (*Plexon Inc*.) with recording sites spaced at 100 µm intervals and referenced to a guide tube on the surface of the brain (Figure 1a). For M1, neural data for the single-exposure visual memory task and categorical single-exposure visual memory task were sometimes recorded from the same electrode penetration. For M2, all neural data were recorded for different electrode penetrations. In both cases the same regions of ITC were targeted. The only difference was whether data was recorded for both tasks from the exact same electrode penetration.

Wideband signals were amplified and digitized at 30 kHz using a Grapevine Data Acquisition System (Ripple Inc.). After high pass filtering of the wideband signal (>300 Hz), using a Butterworth filter, spikes were hand-sorted offline (*Plexon Offline Sorter*) and contain a mixture of single and multi-units. Spike data was quantified as the number of spikes occurring in a given response window. Units were only included in the analysis if they had a significantly greater firing rate after stimulus presentation ([50 350]) compared to before ([-300 0]) as quantified by a t-test with p < 0.05.

LFPs were extracted by low pass filtering wideband signals (<600 Hz) using an 8th-order Type I Chebyshev filter, then downsampling the raw voltage to 1200 Hz, and removing 60 Hz line noise using harmonic regression. ERPs were computed directly from the down sampled and denoised voltage traces. To measure the amplitude of the signal at given frequencies we started by performing a Morlet wavelet decomposition (8 cycles per wavelet) at frequencies ranging from 3 to 500 Hz (produced by linearly sampling frequencies at an interval of 2 Hz) for each trial to estimate the spectrogram. We then converted the amplitude values for each frequency into a percent change from baseline by defining base-line activity as the average amplitude in the window [- 200, 0]. For analyses focused on canonical frequency bands, we defined our ranges as theta (θ): 4-8 Hz, alpha (α): 8-12 Hz, beta (β): 13-30 Hz; gamma (γ): 30-50 Hz; high-gamma activity (HGA): 50-150 Hz, and broad-band: 3-200 Hz. For each trial, activity in a frequency band was quantified as the mean percent change from baseline within a frequency band for a given window computed off the full spectrogram. Following a large literature, we focused LFP analyses primarily on HGA, defined here as (50-150Hz). This range reflects the broadband gamma frequencies commonly observed for HGA in humans and non-human primates. This also limits incorporating divergent spectral effects in lower frequencies and the attenuation in amplitude that occurs with increasing frequency.

Initial exclusion criteria for behavior and spiking data quality are reported in [34] for M3 and M4 and in [46] for M1 and M2 on the single-exposure visual memory task. Given that the field potentials had not previously been analyzed, we took additional measures to assess the quality of the field potentials for each recording session and eliminated additional sessions based on these criteria. Specifically, for each session we plotted the ERP for each channel averaged separately across four consecutive quartiles of trials. Each channel was visually evaluated on two metrics. First, each channel was assessed for instability. Instability was assessed by evaluating whether any two quartiles had noticeable shifts in amplitude, latency, or shape of the ERP. The whole session was eliminated if more than three channels appeared unstable. Second, each channel was assessed for noisy trial-to-trial fluctuations. A channel was eliminated if fluctuations in pre-stimulus voltage varied considerably from the mean or were similar in amplitude to the evoked response and the entire session was again eliminated if more than three channels met this criterion. Importantly, all session elimination decisions were made before conducting any analyses that examined neural representations of experimental variables. Post-hoc analyses revealed that excluded sessions had significantly lower average Pearson correlations between each quartile’s ERP (p=3.4e-5, d’=1.4), capturing the instability and noise across quartiles that formed the basis of session elimination. This led to the elimination of an additional 1 session for M2, and 6 sessions for M3. In addition, two sessions were eliminated because the data files with the raw voltage data could not be analyzed: 1 session for M2 and 1 session for M4. This led to a total of 12 sessions from M1, 7 sessions from M2, 9 sessions from M3, and 9 sessions from M4 included in the analysis.

For the categorical single-exposure visual memory task, data were excluded if there were issues with the experimental set up that rendered the data unusable (1 session for M1, 2 sessions for M2) or there was more than a 1.9x difference in baseline firing rate for a given session between any two blocks (1 session for M1, 2 sessions for M2). No sessions needed to be eliminated due to insufficient data quality of the local field potentials. This led to a total of 17 sessions for M1, and 23 sessions for M2.

### Pseudopopulation alignment

The neural representations in which we are interested require estimates of the responses of hundreds of neurons. Because each session only captures the responses of a few dozen neurons, we align our neural data across sessions into pseudopopulations using the methods reported in [34] and [46] for the single-exposure visual memory task. First, trials were narrowed to only those at the target n-backs (see task details) for which the monkey maintained fixation for the full duration of image presentation and made a behavioral report for both the novel and repeated presentation of an image. This ensures that for every novel image included in the analysis we also have the neural data for its repeated presentation and vice versa. Then, images were grouped by n-back and ordered from lowest to highest memorability images. For the categorical task images were aligned within a block irrespective of their particular n-back but limited to only those in the narrow target range. For sessions that had more images than the session with the smallest number of images in each n-back images were included in the pseudopopulation by subsampling linearly across images ranked for their memorability to include images across the full range of memorabilities. This resulted in pseudopopulations of the following sizes for each monkey in the single-exposure visual memory task: M1-228 images, 440 units; M2-345 images, 281 units; M3-161 images, 227 units; M4 – 103 images, 284 units, and the following sizes for the categorical version of the task: M1-170 images, 692 units; M2-93 images, 648 units.

Once the pseudopopulations were aligned for spiking data, we added the field potentials from the matched channels on which each unit was recorded. When multiple units were sorted off the same channel the field potential data was included multiple times to match the number of units so that comparisons could be made for populations of matched sizes. All cross-validation analyses described below split training and testing by trials (not channels) and as such, repeating data in this way does not result in spill over between training and testing sets. This process yielded pseudopopulations with simultaneously recorded spike and LFP data from matched images on their novel and repeated presentations. All analyses reported in the paper utilized these pseudopopulations.

### Visually evoked responses

PSTHs (Figure 1, Supplemental Figure S1) were computed with a 50 ms window taking 10 ms steps in the range [-200, 500] for M1 and M2 and [-200, 400] for M3 and M4. This smoothing was applied in all cases except the ERP where voltage traces were plotted directly. Grand mean firing rate for spiking activity (Figure 1B) and ERPs (Figure 1C) were baseline subtracted per trial relative to the [-200, 0] window. Spectrograms were first computed as an average across images for each channel and then averaged across channels (Figure 1D). Error shadows show 95% confidence intervals arrived at by 1000 iterations of bootstrapped resampling across units (spikes) or channels (LFPs) with replacement.

### Novelty signals

PSTHs broken out by novel and repeated images (Figure 2A-B, Supplemental Figure S5) and their error shadows were computed in the same way as Figure 1 but separately for novel and repeated images. The repetition effect (Figure 2C-D, Figure 5B) was computed with the following equation:

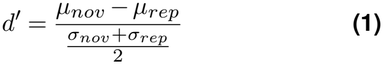

where *µ*_*nov*_, *µ*_*rep*_, *σ*_*nov*_, and *σ*_*rep*_ are the mean and standard deviation of responses to novel and repeated images respectively across images. This was computed per unit or channel (Figure 2C, Figure 5B) or for the average response across units or channels (Figure 2D). Positive values correspond to repetition suppression and negative values to repetition enhancement.

The cross-validated decoder (Figure 2E, Figure 5C) was previously reported in [34] (Spike Count Classifier). Briefly, the population response **x** was quantified as the vector of spike counts for a given pseudoimage. The decoder took the form of a linear decoding axis:

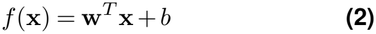

where **w** is an N-dimensional vector (and N is the number of units) containing the linear weights applied to each unit. The weight applied to each unit was 1/N and *b* was calculated as:

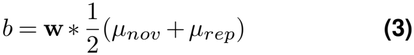

The classifier was trained separately for each neural measure and evaluated in a cross-validated manner, with each of 50 iterations training on a random 80% of images and testing on the remaining 20%. On each iteration the responses of each unit were randomly shuffled to ensure that artificial correlations (e.g., between the units recorded in different sessions) were removed. To evaluate performance as a function of population size, on each iteration of cross-validation **x** was composed of a randomly subsampled number of units or channels of a given population size. Error bars show the standard deviation of performance across 50 iterations of subsampling and cross-validation (Figure 2F).

To compare classifier performance growth as a function of population size, the relationship between performance and population size was fit separately for spikes and HGA for each monkey using Matlab’s `fit()` function with the following power law function:

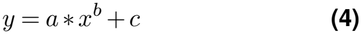

where *y* is the performance of the classifier, *x* is the population size, and *a, b*, and *c* are parameters fit separately for each model. We then used these fits to estimate the population size required to reach 75% classifier performance and compared population sizes needed for spikes and HGA within each monkey. This yielded the following ratios for each monkey: 5.2x for M1, 3.6x for M2, 7.1x for M3, and 3.7x for M4. We conservatively take the lowest of these numbers in claiming that it requires at least 3-fold less data to reach matched decoding performance with HGA as compared to spikes.

### Memorability

We obtained memorability scores using the procedure documented in [78]. Briefly, we used MemNet [80], a convolutional neural network trained to estimate image memorability on a large-scale dataset of natural images, to estimate image memorability scores as would have been measured from human performance on a memory task. Scores can range from 0 (least memorable) to 1 (most memorable) and achieve a rank correlation of 0.64 with mean human-estimated memorability, close to the upper bound on the consistency between human-based scores which has a rank correlation of 0.68. For neural analyses (Figure 3D-E), the memorability score reported for a pseudoimage is the mean memorability across the images aligned to create that pseudoimage, and population responses are the mean across all units or channels averaged across novel and repeated presentations responding to the presentation of the images that make up that pseudoimage.

### Neural predictions of behavioral performance

Neural predictions of behavioral performance were estimated using a Fisher Linear Discriminant (FLD) decoder and rescaled to minimize mean squared error (MSE) as described in [34]. The FLD is implemented according to Equations 2 and 3, with the weights determined by the following equation:

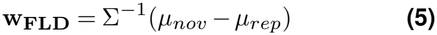

Where Σ^*−*1^ is the inverse of the covariance matrix averaged across novel and repeated presentations. As previously reported [34, 44–46], due to the large amounts of data required to estimate the off-diagonal elements of the covariance matrix, we set all off-diagonal terms to 0 and consider only the diagonal of the covariance matrix. Intuitively, this means that the resulting classifier essentially weights each unit by its d prime.

The FLD was trained in the same cross-validated manner as the Spike Count Classifier, but with 2 images per n-back held out for cross-validation over 5000 iterations rather than the 80/20 split used for the Spike Count Classifier across 50 iterations. Here, the FLD was used as opposed to the Spike Count Classifier because it was previously shown to be a better match to behavior and because it allows for differential weighting of individual units or channels with either positive or negative weights (repetition suppression or enhancement respectively). This gave each neural measure its “best shot” at estimating performance without enforcing that all units had to show repetition suppression. Behavioral predictions were made using neural data averaged between 200 and 500 ms (M1/M2) and 200 and 400 ms (M3/M4) after stimulus onset because this analysis window gave slightly better behavioral predictions across most frequency bands.

Classifier performance was rescaled to provide minimum MSE with behavior using the methods described in [34] to estimate performance for larger-sized populations. Briefly, following on previous observations that the trajectories of means of the FLD scale linearly with increasing population size while the standard deviations plateau, we separately established for each neural measure the rate of growth of the mean of the FLD and peak standard deviation as a function of population size. We used a linear extrapolation from the two largest population sizes to estimate how the means at each n-back would grow as a function of larger population sizes, set the standard deviation to the plateaued value, and evaluated performance for these larger population sizes under the assumption that the neural responses are Gaussian as parameterized by these mean and standard deviation values. The classifier performance for each neural measure was selected as the population size that produced the best match to behavior as measured by MSE. We emphasize that this rescaling procedure increases the level of classifier performance without changing the general trends of performance with n-back. This method allows us to find the best possible fit to behavior for each neural measure making comparisons between them fair and eliminating any confounding factor of the amount of data needed to reach classifier performance that matches behavior.

The prediction quality (PQ) of each classifier was defined by the following equation:

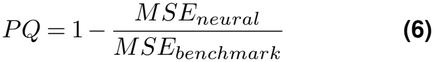

where *MSE*_*neural*_ is the mean-squared error (MSE) between the neural prediction and behavior and *MSE*_*benchmark*_ is the MSE between behavior and a “worst-case scenario” benchmark wherein performance is at 50% across all n-backs. PQ values are bounded at 1 if the neural predictions are a perfect match to actual behavior while PQ values of 0 correspond to chance performance across all n-backs. PQ values can go below 0 if performance dips below 50%.

### Quantifying categorical representations

All analyses of categorical representations (Figure 5D-E) utilized data collected during the categorical single-exposure visual memory task. Representational similarity matrices (Figure 5D) were computed by first z-scoring each unit (spikes) or channel’s (HGA) average activity across novel and repeated presentations of each image and then correlating the vector of z-scored responses across all images for all images in the pseudopopulation.

The cross-validated prototype categorical decoder (Figure 5E) performs a five-way classification to determine which category a given response vector belongs to. On each iteration of cross-validation, one image from each category is randomly selected as a “test” image. The response of each unit (spikes) or channel (HGA) to all other images in each category is averaged to produce a “prototype” vector for each category. Each of the test images is then correlated with the five prototypes and is classified as belonging to the category with whose prototype it correlates most strongly. Figure 5E shows the result of applying this decoder to the pseudopopulation with 1000 iterations of cross-validation.

## Supplementary Information

**Figure S1.**
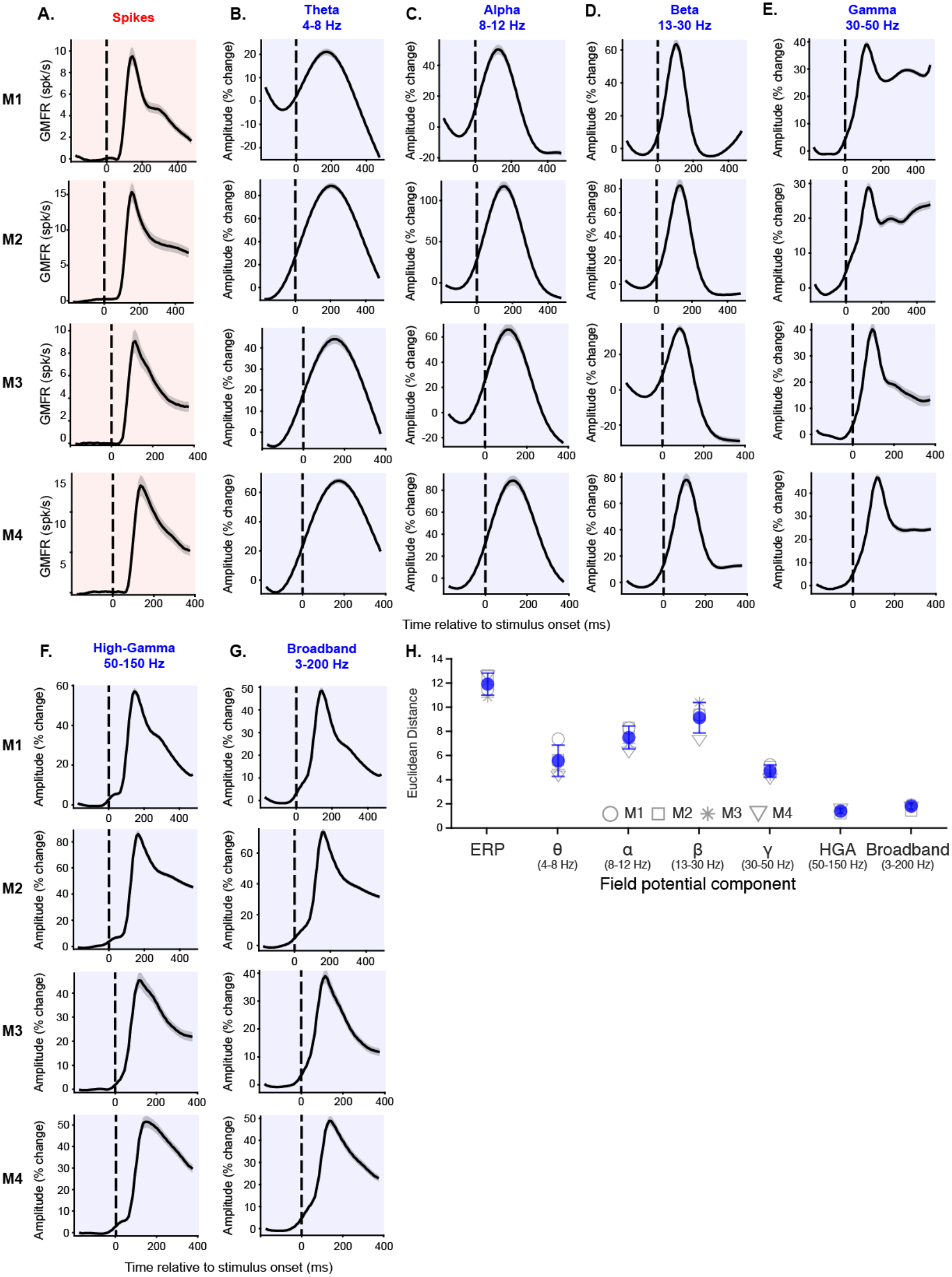
HGA captures population spiking vigor better than other features of the LFP. Each row is the neural data from one monkey. Spikes are shaded in red and features of the LFP in blue. Stimulus onset is marked with the dashed vertical black line. All measures are baseline subtracted with respect to activity in the window [-200, 0] and shown as a function of time relative to stimulus onset. Error shadows are bootstrapped 95% confidence intervals (see Methods). **A**. Grand mean firing rate. **B**. Amplitude of activity in the theta band (4-8 Hz). **C**. Amplitude of activity in the alpha band (8-12 Hz). **D**. Amplitude of activity in the beta band (13-30 Hz). **E**. Amplitude of activity in the gamma band (30-50 Hz). **F**. Amplitude of activity in the high-gamma band (50-150 Hz). **G**. Amplitude of broadband activity (3-200 Hz). **H**. Euclidean distance between the z-scored event-related time course of activity for each component of the LFP compared to spiking activity. Each shape is an individual monkey, the blue dots depict the mean across the four monkeys, and the error bars depict the standard deviation.

**Figure S2.**
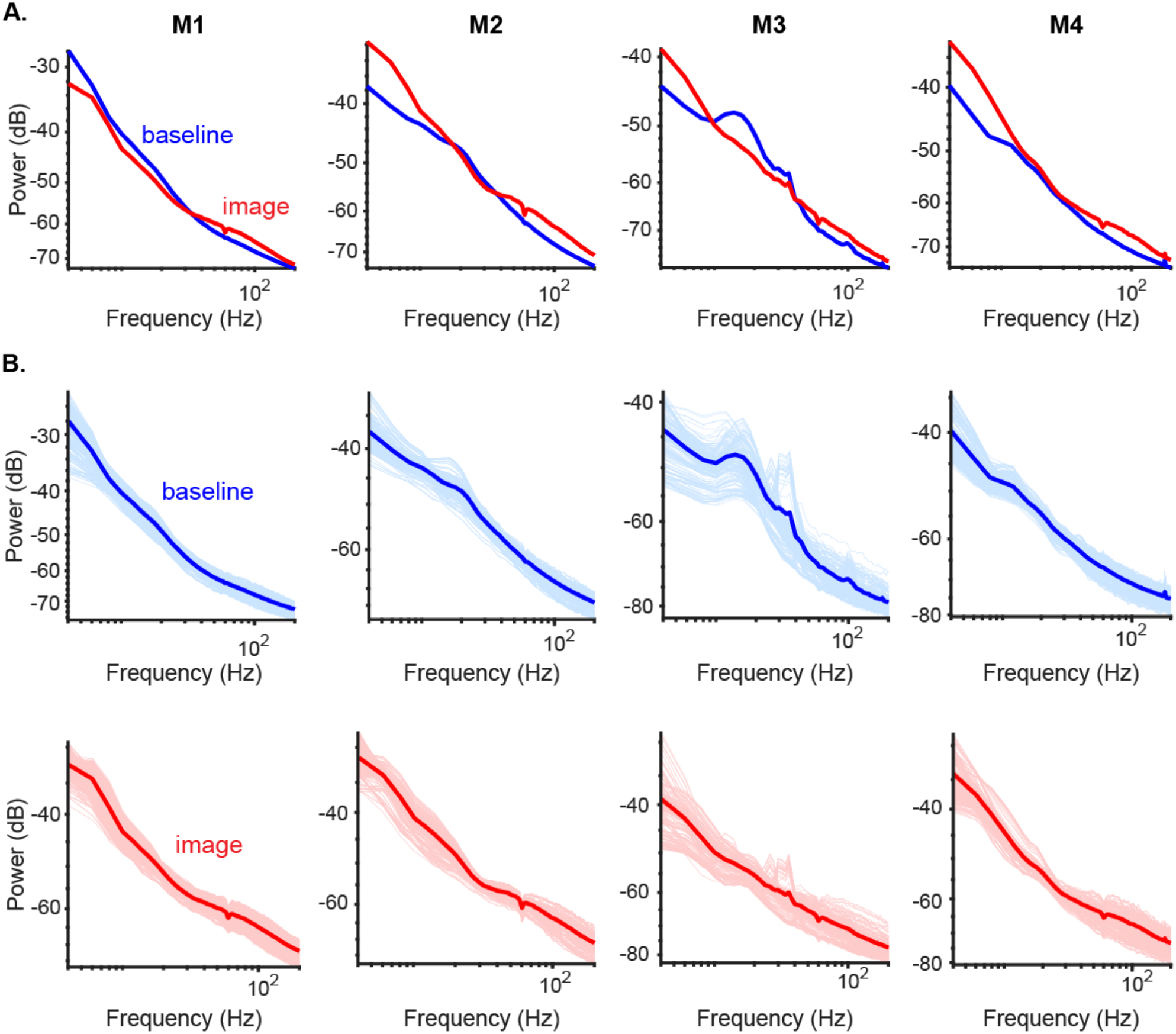
The power spectra for all four monkeys show no consistent evidence of oscillatory peaks. **A**. The power spectrum for each monkey averaged across all recorded channels for the baseline (blue, [-500 0] for M1 and M2, [-400 0] for M3 and M4) and image (red, [0 500] for M1 and M2, [0 400] for M3 and M4) periods. The spectra show evidence of a clear broadband power increase following stimulus presentation but limited and inconsistent stimulus-driven oscillatory peaks. **B**. Each column is the data from one monkey. The top row (blue) shows the average power spectrum across channels (dark blue) and the spectra from individual channels (light blue) in the baseline period. The bottom row shows the average power spectrum across channels (dark red) and the spectra from individual channels (light red) in the image period. The shape of the power spectrum with no clear oscillatory bumps is consistent across most channels, with some evidence for oscillatory activity in M3’s baseline activity and a few channels of M3’s activity in the image period, but not consistently across monkeys. To quantify this observation, we used Matlab’s findpeaks() function to identify putative peaks on each channel during the image period. This analysis identified 0% of channels in M1 and M2, 18.1% of channels in M3 and 8.1% of channels in M4 as having a peak using a liberal threshold of a minimum 3 dB prominence (height from trough to putative peak) and 2 Hz width.

**Figure S3.**
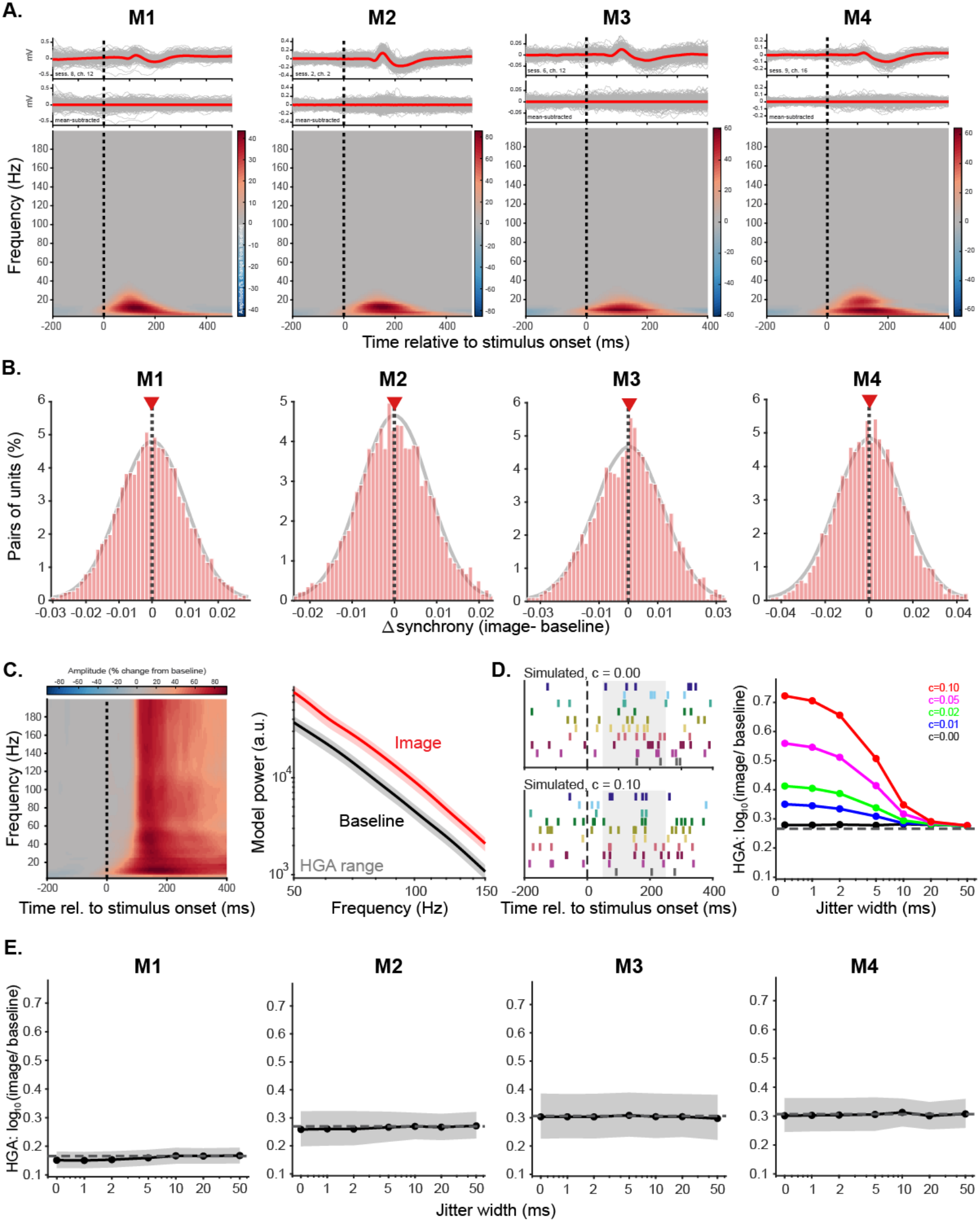
HGA in our recordings largely reflects asynchronous neural activity. To investigate the extent to which synchronous versus asynchronous activity contributes to the HGA observed in our dataset we considered two possible synchronous contributors to HGA: ERPs (**A**) and synchronous neural firing (**B-E**). While both could contribute to HGA, in theory, our analyses suggest HGA in our dataset does not reflect either of these contributors and is instead largely reflected by asynchronous neural activity. **A**. Contribution of the ERP to spectral power shown separately for each monkey. To identify the contribution of large, synchronized voltage changes (ERPs) following image presentation to HGA, we recomputed the spectrograms after subtracting the ERP from each trial separately for each channel. Top: Voltage responses from one example channel derived from the raw voltage. Individual trials are shown in gray with the ERP overlaid in red. Middle: Voltage responses from the same example channel shown on top after mean subtracting the raw voltage. Individual trials are shown in gray with the mean overlaid in red. Note that the mean response is zero by construction, but variability across trials is still preserved. Bottom: The difference between the spectrograms computed from the raw and mean-subtracted ERPs. This difference shows the relative contribution of the ERP to spectral power. To quantify the contribution of the ERP to HGA, we computed the ratio of the mean HGA in the difference spectrogram (Supplemental Figure S3A) to the raw spectrogram (Figure 1D). We found that the ERP accounted for 0.0-0.6% of HGA across monkeys (95% CIs bootstrapped across units ranging from 0.0% to 0.7% across monkeys). This demonstrates that the ERP was reflected strongly in spectral power, but only at lower frequencies well below the HGA frequency range. **B**. Distribution of the change in synchrony following image presentation for all pairs of units simultaneously recorded on different and non-adjacent channels shown separately for each monkey. The synchrony index was computed by measuring the excess number of coincident spikes within a +/-2 ms window. Specifically, we counted the number of raw coincidences in this window, subtracted the expected number of coincidences due to firing rate alone (by computing coincidences when spike times occur at a random time within a 25 ms interval), and then divided by the geometric mean of the two units’ spike counts to normalize for firing rates across all pairs. The synchrony index is the difference between this measure during the image presentation ([50 250]) and baseline ([-200 0]) periods (i.e., a measure of whether firing rate changes following image presentation are accompanied by changes in neural synchrony). The distribution of this synchrony index across all pairs is shown by the red bars with the means depicted by the solid red triangles. The black dashed line depicts a difference of 0. A null distribution was computed by randomly jittering spike times within a 25 ms window (separately for each period), recomputing the synchrony index in an identical manner across 200 iterations, and then pooling all iterations across all pairs. The distribution was smoothed and plotted as the solid gray line. The distributions of synchrony indices observed in our data were indistinguishable from the null distribution (p=0.11-0.60 across monkeys, where we computed p as the fraction of 200 randomly shuffled spike times whose mean synchrony index was at least as large as the observed mean). **C**. To test the contribution of neural synchrony in our population to HGA we implemented the model proposed by [14] to predict an LFP for each session directly from our spiking activity. Specifically, we convolved each unit’s spiking activity with an exponential decay filter (Tau=2.3 ms) and then summed the convolved activity across all units and passed it through a leaky integrator (100 ms) and a 0.5 Hz high-pass filter (much like a recording system) to predict an LFP for each session. Left: The spectrogram computed from the modeled LFPs and averaged across sessions for one example animal (M4). The predicted power spectrum captures the strong broadband changes observed following stimulus onset, including in the HGA frequency range. Right: A snapshot of the power in the HGA frequency range (50-150 Hz) during the baseline (black, [-200 0]) and image presentation (red, [50 250]) periods for the same monkey. Shaded error bars show the bootstrapped 95% confidence interval across all sessions. The predicted power spectra demonstrate the key hallmark of [14], being a broadband shift upward with increasing firing rate. **D**. To quantify to what extent neural synchrony contributed to the observed HGA we jittered the spike times in our neural data in windows with increasing width (x axis) and then passed them through our implementation of the [14] model to look at how the ratio of HGA in the image presentation and baseline periods of the modeled LFP changed. The logic is that if neural synchrony is contributing to evoked HGA then jittering spike times to remove the synchrony should decrease the HGA measured in the image presentation period. First, to demonstrate that this analysis is capable of measuring an impact of neural synchrony on HGA we ran a positive control analysis where we simulated spike trains with varying levels of synchrony and ran them through the exact same proposed jitter analysis pipeline. We found that the pipeline was indeed sensitive enough to detect even small levels of neural synchrony induced by image presentation. Left: Rasters showing 10 example spike trains simulated with no synchrony (top) and strong synchrony (bottom). Specifically, to simulate spike trains in the image presentation period with various levels of synchrony we adopted the multiple interaction process method from [81]. We began by generating one spike train with a high firing rate to seed the simulated spike trains. Spike trains to be included in the population were then generated by copying a proportion c of the spikes from the original train independently 30 times to simulate 30 units. This was repeated 300 times to simulate 300 trials. With a higher level of c, spike times are more likely to be identical, which increases neural synchrony (top v. bottom). Right: The ratio of HGA measured in the image v. baseline period as a function of the window size used to jitter spike times. Each colored line is the result of running this analysis on a simulated population with a different level of neural synchrony. Even with small levels of synchrony (c=0.01) the analysis detects a drop in the observed ratio as jitter increases. The grey dashed line shows the ratio predicted by comparing firing rate in each period alone. A population with no synchrony maintains this ratio regardless of jitter (black line), while populations with some neural synchrony converge toward this ratio as jitter increases and the influence of synchrony on image presentation period HGA is removed (colored lines). **E**. The same plots as in D after running the jitter analysis on the real neural data, shown separately for each monkey. Y axes were selected to match those in the simulation in D for ease of comparison. Once again, the grey dashed line shows the ratio based on firing rates in the baseline and image periods. Shaded error bars are the bootstrapped 95% confidence interval across sessions. All four monkeys show patterns most similar to the simulated population with no neural synchrony induced by image presentation.

**Figure S4.**
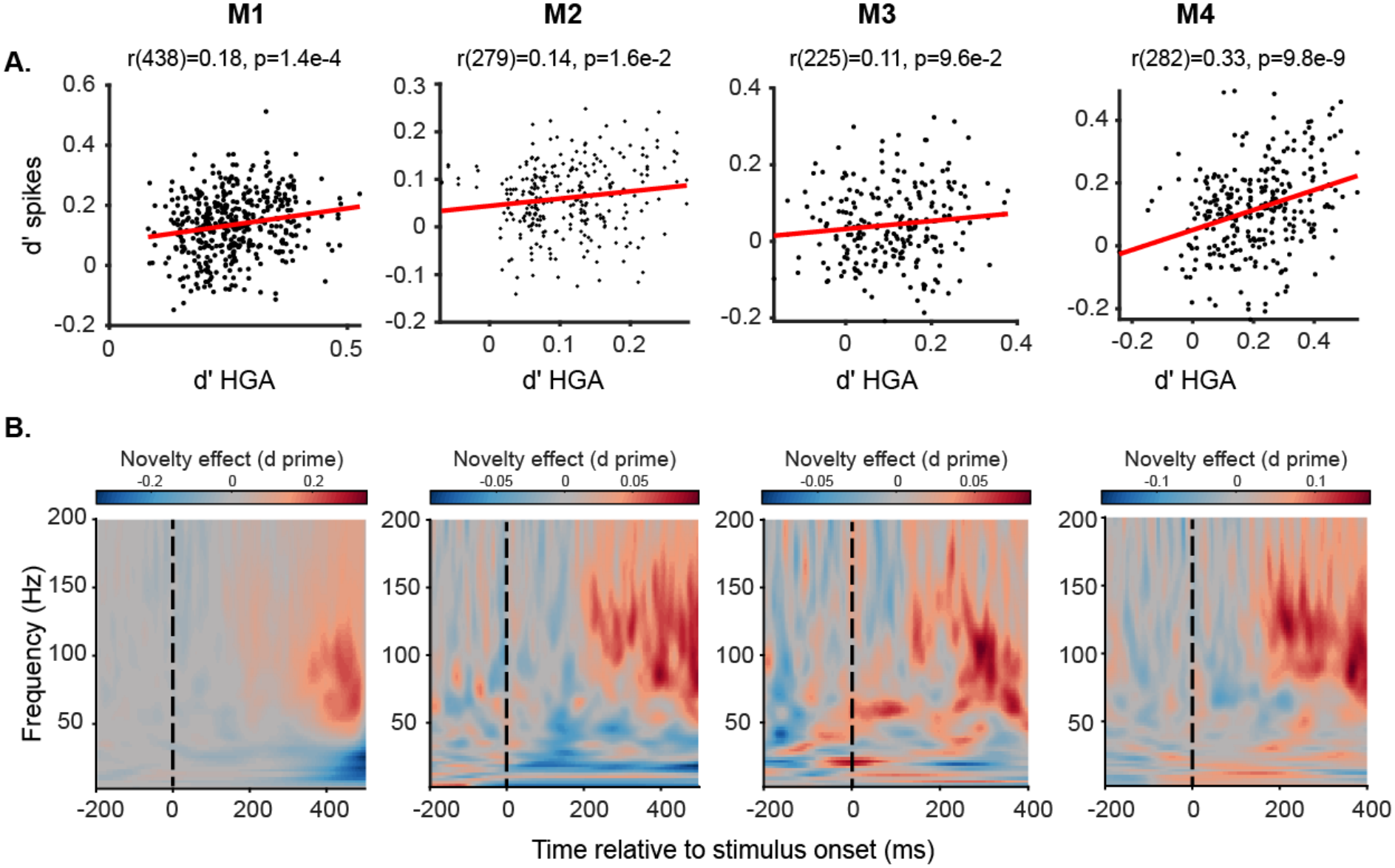
Repetition suppression on matched channels and across the full frequency spectrum. **A. Spikes and HGA recorded on the same channel are weakly positively correlated**. Pearson’s correlation between the d prime between the response to novel and repeated presentations of an image of a given unit and its corresponding channel’s HGA for each monkey. **B. Repetition suppression evolves late and only at higher frequencies**. Full spectrogram of d prime between the response to novel and repeated images. D prime was computed from the baseline-subtracted spectrogram of response to novel and repeated images for each channel and then averaged across channels.

**Figure S5.**
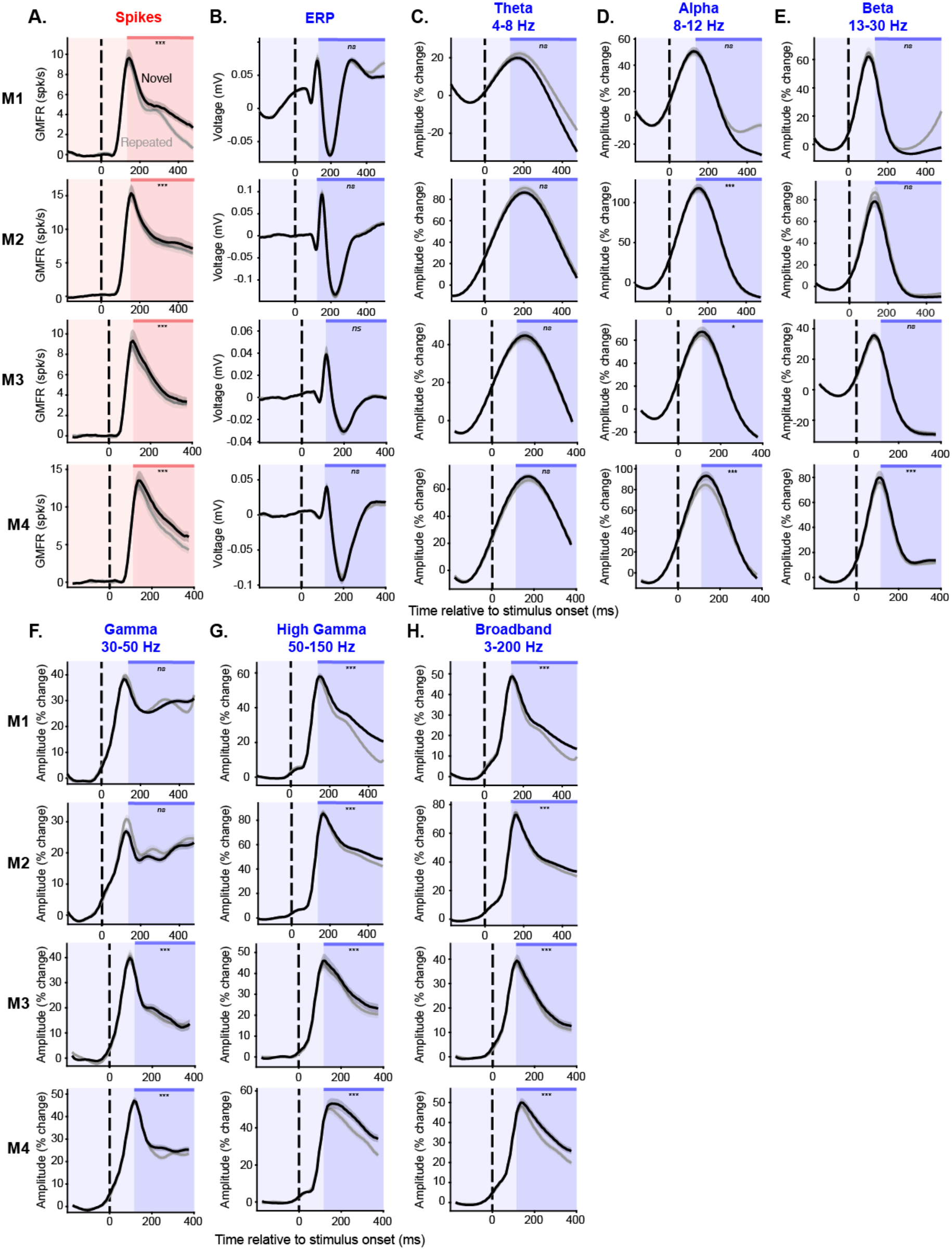
HGA captures repetition suppression better and more consistently than any other features of the LFP. Each row is the neural data from one monkey. Stimulus onset is marked with the dashed vertical black line. All measures are baseline subtracted with respect to activity in the window [-200, 0] and shown as a function of time relative to stimulus onset. Error shadows are bootstrapped 95% confidence intervals (see Methods). Significance tests are for a one-tailed Wilcoxon signed rank test (novel > repeated). ns: non-significant, *: p<0.01, **: p<0.001, ***: p<0.0001. Shaded regions show the window used for significance testing (150 ms to the end of stimulus presentation). **A**. Grand mean firing rate. **B**. Magnitude of voltage fluctuation (ERP). **C**. Amplitude of activity in the theta band (4-8 Hz). **D**. Amplitude of activity in the alpha band (8-12 Hz). **E**. Amplitude of activity in the beta band (13-30 Hz). **F**. Amplitude of activity in the gamma band (30-50 Hz). **G**. Amplitude of activity in the high-gamma band (50-150 Hz). **H**. Amplitude of broadband activity (3-200 Hz).

**Figure S6.**
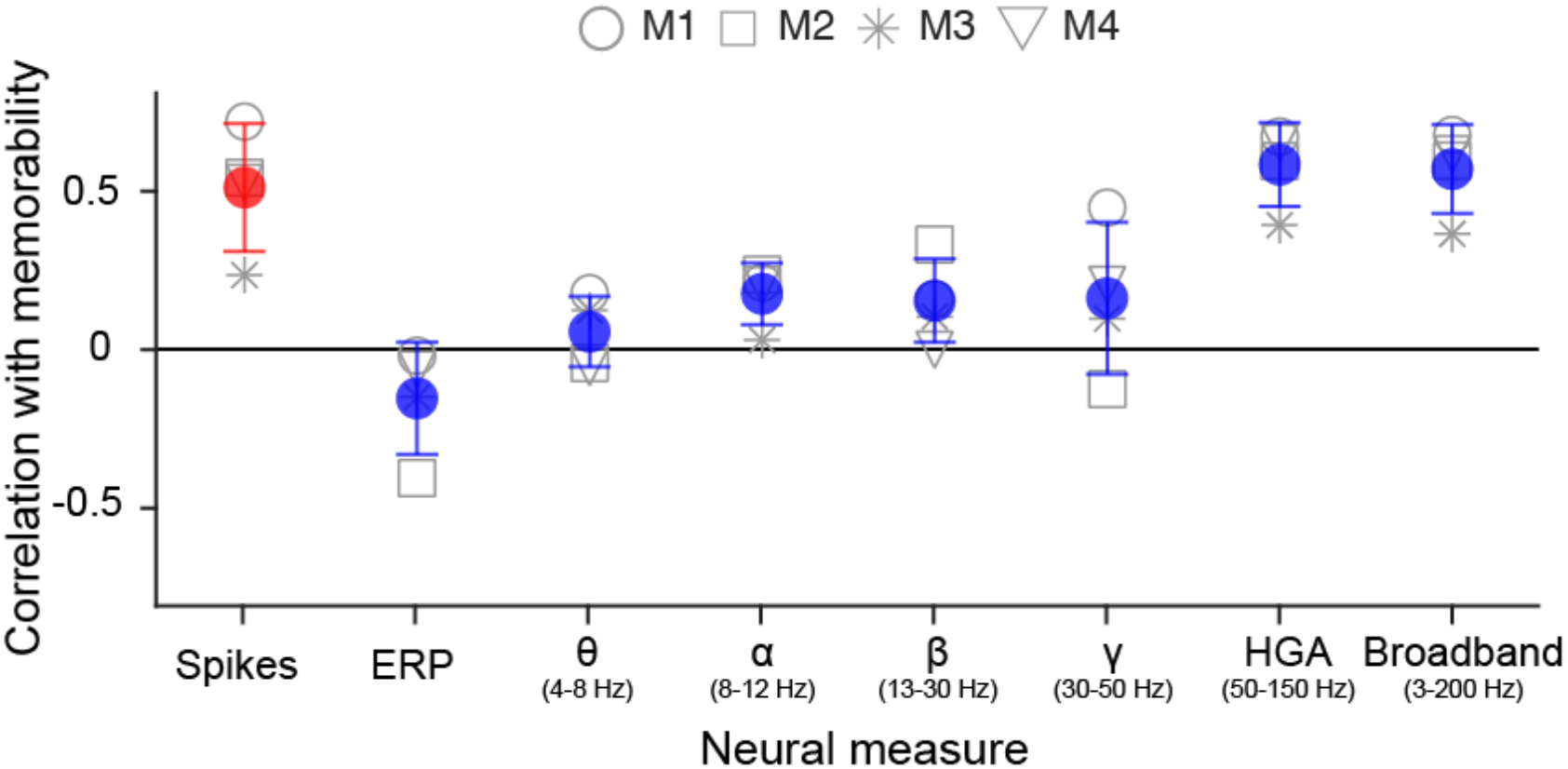
The correlation between memorability and spiking activity is only captured by high frequency components of the LFP (not low frequencies). Correlation between memorability score and response magnitude averaged over all channels for spikes (red) and several features of the LFP (blue). As was observed in novelty signals, the memorability correlation observed in spiking activity is only achieved in high frequencies of the field potential. Each point is a single monkey, colored points are the average across monkeys, and error bars are the standard deviation across all four monkeys.

**Figure S7.**
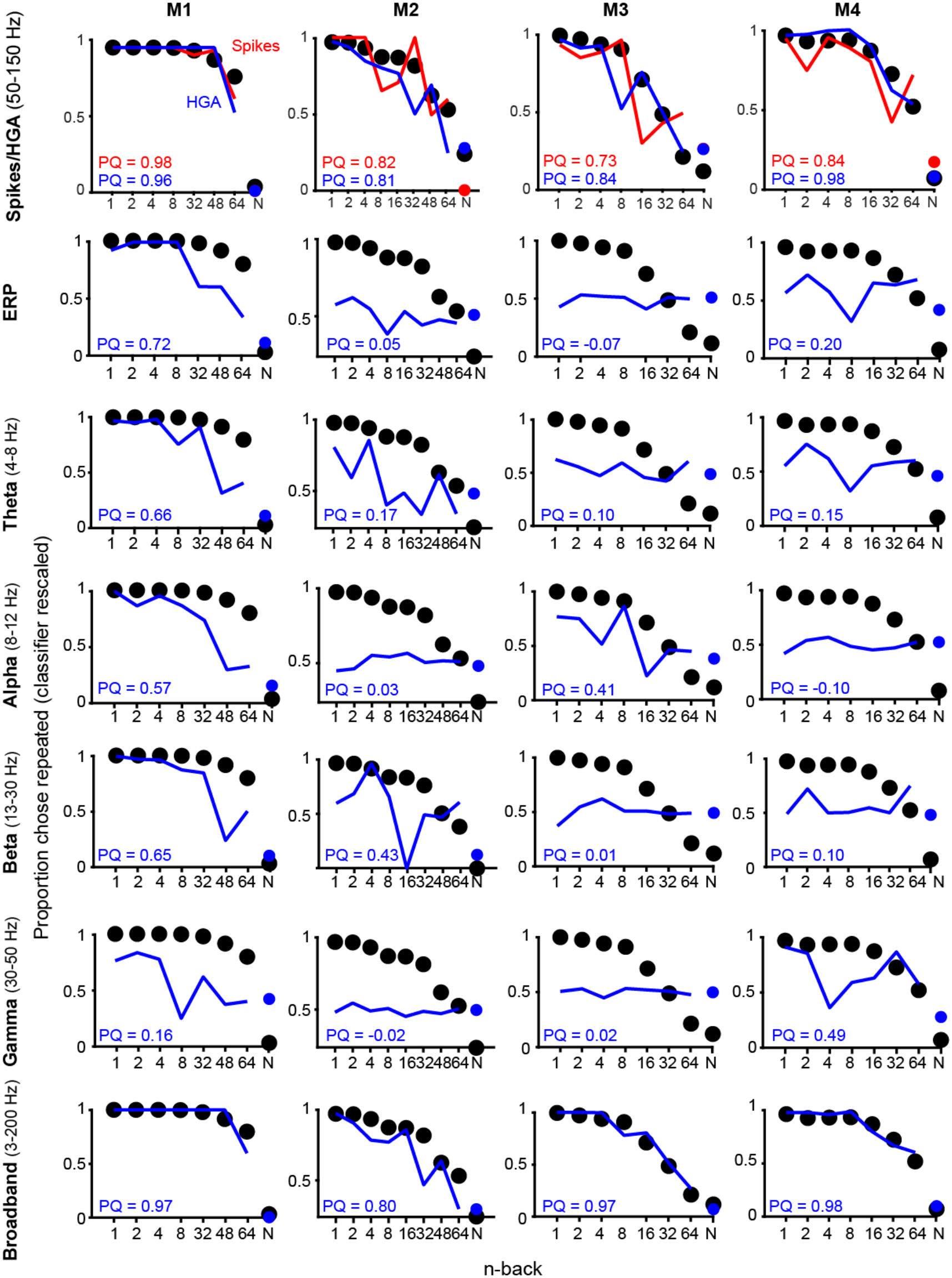
HGA, but not lower frequency components, predicts forgetting behavior as well as spikes. Neural predictions of the Fisher Linear Discriminant classifier (see Methods) trained on spikes (red) and features of the LFP (blue) rescaled to match the level of behavioral performance (black) for all four monkeys (see Methods). Each row shows results from a different frequency band. Prediction Quality (PQ) values measure how well classifier performance predicts behavior (see Methods). The x-axis denotes the n-back for repeated images and the average performance for novel images is labeled as “N”.

**Figure S8.**
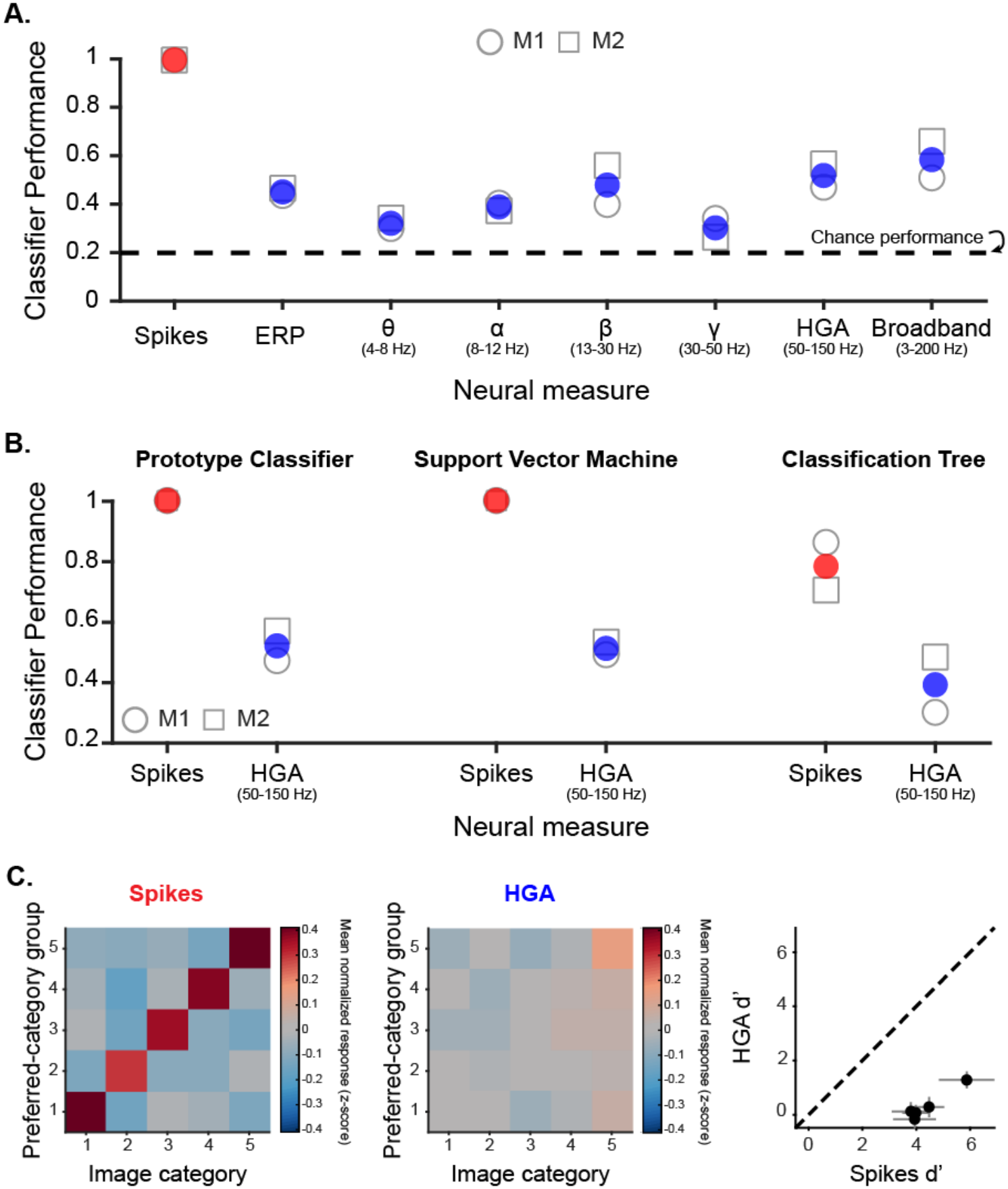
Weaker categorical decoding compared to spikes is unique to HGA and cannot be accounted for with different classifiers or by grouping channels by tuning of spiking activity. **A**. Prototype category decoder shown in Fig. 5 with performance broken out as a function of frequency band. All frequency bands show worse decoding performance than spikes. Unlike the visual memory signals previously reported (novelty, recency, memorability), HGA is no closer to spikes than any other frequency band. **B**. Performance of three types of categorical decoders in performing a 5-way image category classification: Prototype classifier (left, reported in Fig. 5), support vector machine (middle), and classification tree (right). Chance is 0.2 for all decoders. Each shape represents classifier performance when trained on a single monkey and the colored circles are the mean across the two monkeys for spikes (red) and HGA (blue). For all three classifiers, performance is higher for spikes than HGA, indicating stronger categorical representations in spikes than HGA. **C**. To validate that the misalignment between spikes and HGA for categorical representations is indeed due to a salt and pepper anatomical organization of similarly tuned neurons we explicitly sorted units post-hoc based on their categorical tuning and asked if these functionally defined clusters might show better alignment between spikes and HGA. We started by categorizing each unit based on which of the five object categories it responded to most. To avoid circularity, we did this in a cross-validated manner, using a training set of images to identify the unit’s category selectivity and visualizing its firing rate using a held-out test set of images across 100 iterations of cross-validation. Plots show results of combined data across the two monkeys participating in the categorical single exposure visual memory task (M1 and M2). **Left**: Average z-scored spiking response of units with a given category preference to images in each category. The diagonal structure indicates that the cross-validated category tuning procedure accurately captures underlying tuning preferences. **Middle**: Average z-scored response of HGA recorded on matched channels to the units sorted and shown on the left. That there is no diagonal like the spiking activity indicates that the category tuning present when clustering spiking activity is not present in the simultaneously recorded HGA. **Right**: On each split we computed the d-prime between the responses to images from the preferred category compared to images from all other categories for all five categories. Each point, corresponding to one of the five categories, shows the mean d-prime for the spikes and HGA across splits. Points below the diagonal showed stronger category tuning in spikes than HGA. Error bars depict the standard deviation across splits.

**Figure S9.**
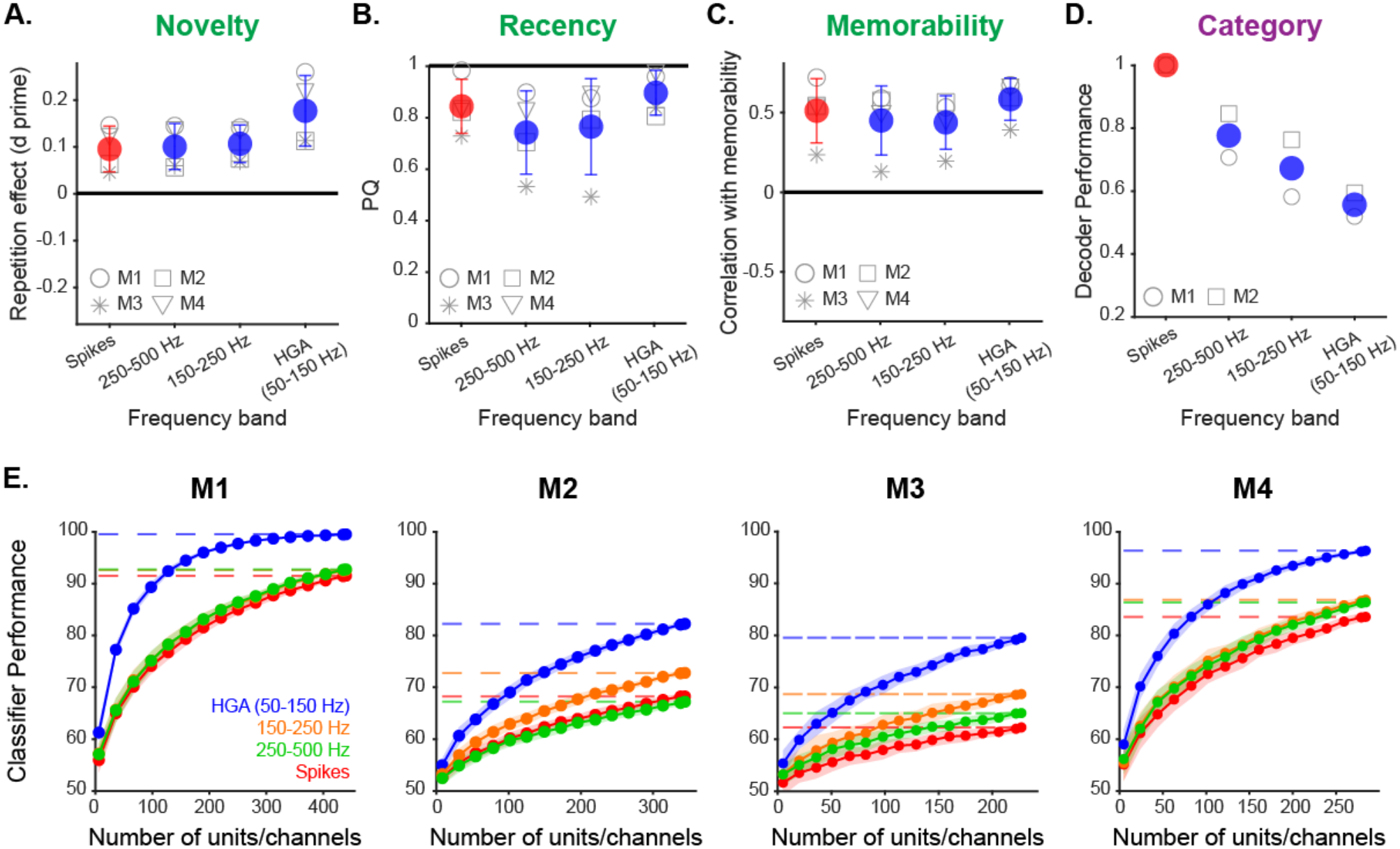
Representations in higher frequency activity of the LFP (> 150 Hz) converge toward spiking representations. **A-D**. Various neural representations compared between spikes (red) and three frequency bands (blue). Each shape corresponds to the value for a single monkey and the colored circles show the mean across monkeys with error bars depicting the standard deviation across monkeys. Error bars are omitted from D which had only two monkeys. For magnitude codes (green) representations in higher frequency bands are comparable to spikes. For the pattern-of-spikes code, category, neural representations converged toward spikes as frequency bands got higher. **A**. Repetition effect, as measured as d prime between the response to novel and repeated presentations of images for each unit or channel and then averaged across units/channels. **B**. Prediction quality of a classifier trained to distinguish novel from repeated images in predicting memory as a function of image recency (see Methods). **C**. The correlation between the mean neural response across the population and image memorability. **D**. Performance of a prototype classifier trained to predict the category of a given image (see Methods). **E**. Performance of a spike count classifier trained to distinguish novel from repeated images on the basis of response magnitude as a function of the number of units or channels included in the analysis (see Methods). While HGA, which is thought to aggregate signals over a larger volume of tissue than higher frequencies [48, 63], increases performance faster than spikes, higher frequency bands converge toward spikes, likely reflecting the smaller volume of tissue over which they aggregate activity.

## Supplementary Tables

**Table S1.**
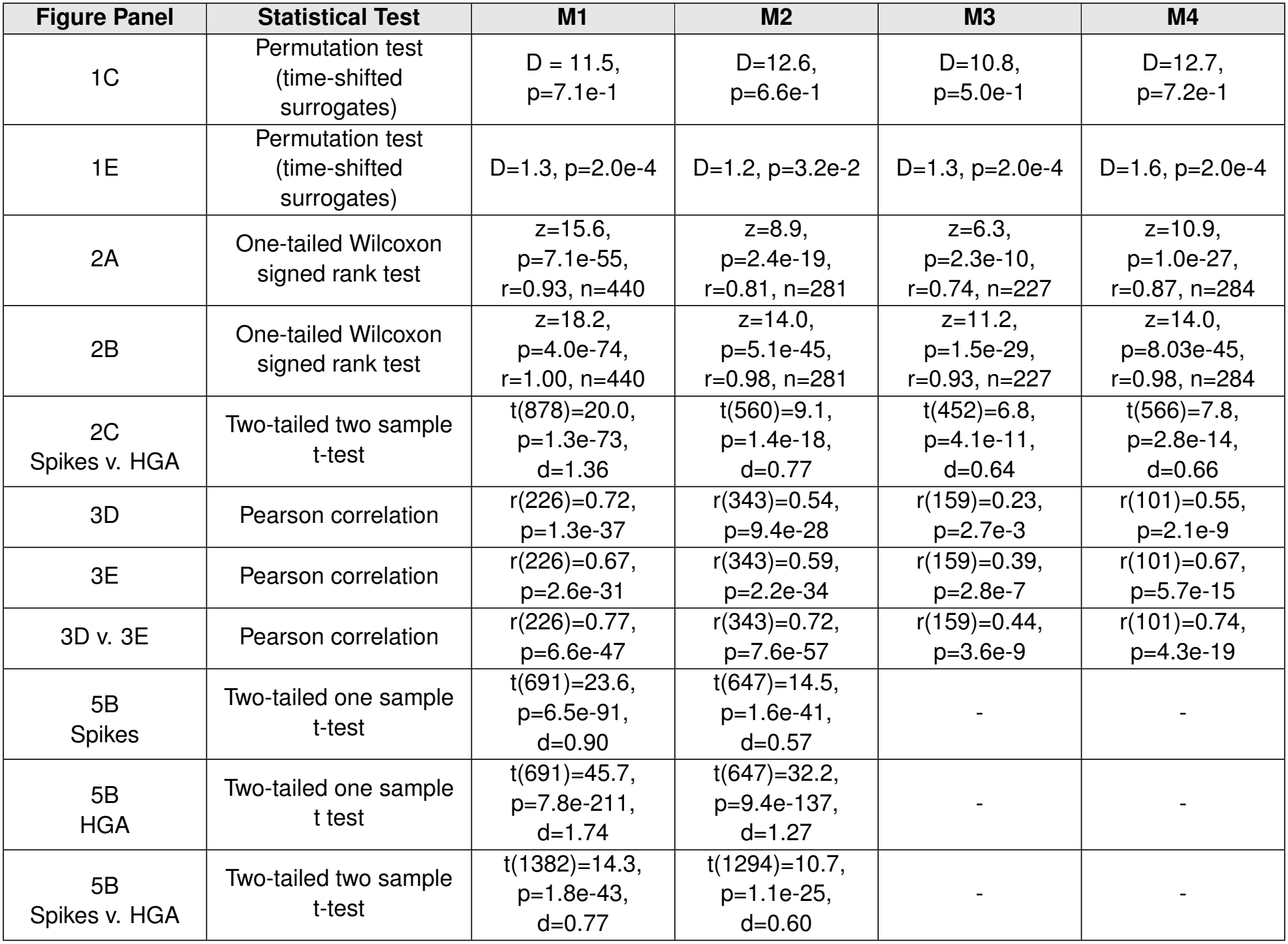
Statistical Tests. This table contains results of the statistical tests reported throughout the paper. **For 1C and 1E**: Significance of the Euclidean distance between the normalized stimulus-evoked responses was assessed by comparing the observed Euclidean distance to a null distribution produced by randomly shifting the response and computing Euclidean distance between the shifted responses across 5000 permutations. D is the observed Euclidean distance between the z-scored stimulus-evoked responses and p is the proportion of shuffles that produced distances at least as similar as the observed distance. **For 2A and 2B**: Effect size, r, is computed as the sum of ranks of positive differences (novel > repeated) divided by the sum of all ranks. Therefore r = 0.5 corresponds to no difference in the response to novel and repeated images across all units/channels and r=1 corresponds to every unit or channel having a positive shift.

| **Figure Panel** | **Statistical Test** | **M1** | **M2** | **M3** | **M4** |
|---|---|---|---|---|---|
| 1C | Permutation test (time-shifted surrogates) | D = 11.5, p=7.1e-1 | D=12.6, p=6.6e-1 | D=10.8, p=5.0e-1 | D=12.7, p=7.2e-1 |
| 1E | Permutation test (time-shifted surrogates) | D=1.3, p=2.0e-4 | D=1.2, p=3.2e-2 | D=1.3, p=2.0e-4 | D=1.6, p=2.0e-4 |
| 2A | One-tailed Wilcoxon signed rank test | z=15.6, p=7.1e-55, r=0.93, n=440 | z=8.9, p=2.4e-19, r=0.81, n=281 | z=6.3, p=2.3e-10, r=0.74, n=227 | z=10.9, p=1.0e-27, r=0.87, n=284 |
| 2B | One-tailed Wilcoxon signed rank test | z=18.2, p=4.0e-74, r=1.00, n=440 | z=14.0, p=5.1e-45, r=0.98, n=281 | z=11.2, p=1.5e-29, r=0.93, n=227 | z=14.0, p=8.03e-45, r=0.98, n=284 |
| 2C<br>Spikes v. HGA | Two-tailed two sample t-test | t(878)=20.0, p=1.3e-73, d=1.36 | t(560)=9.1, p=1.4e-18, d=0.77 | t(452)=6.8, p=4.1e-11, d=0.64 | t(566)=7.8, p=2.8e-14, d=0.66 |
| 3D | Pearson correlation | r(226)=0.72, p=1.3e-37 | r(343)=0.54, p=9.4e-28 | r(159)=0.23, p=2.7e-3 | r(101)=0.55, p=2.1e-9 |
| 3E | Pearson correlation | r(226)=0.67, p=2.6e-31 | r(343)=0.59, p=2.2e-34 | r(159)=0.39, p=2.8e-7 | r(101)=0.67, p=5.7e-15 |
| 3D v. 3E | Pearson correlation | r(226)=0.77, p=6.6e-47 | r(343)=0.72, p=7.6e-57 | r(159)=0.44, p=3.6e-9 | r(101)=0.74, p=4.3e-19 |
| 5B<br>Spikes | Two-tailed one sample t-test | t(691)=23.6, p=6.5e-91, d=0.90 | t(647)=14.5, p=1.6e-41, d=0.57 | - | - |
| 5B<br>HGA | Two-tailed one sample t test | t(691)=45.7, p=7.8e-211, d=1.74 | t(647)=32.2, p=9.4e-137, d=1.27 | - | - |
| 5B<br>Spikes v. HGA | Two-tailed two sample t-test | t(1382)=14.3, p=1.8e-43, d=0.77 | t(1294)=10.7, p=1.1e-25, d=0.60 | - | - |

## Notes

### Competing Interest Statement

The authors have declared no competing interest.

### Summary of Updates

Further analyses exploring the nature of the neural spiking activity that is reflected in high-gamma activity (HGA)

